# Apicobasal surfaceome architecture encodes for polarized epithelial functionality and depends on tumor suppressor PTEN

**DOI:** 10.1101/2020.11.02.365320

**Authors:** Anika Koetemann, Bernd Wollscheid

## Abstract

The loss of apicobasal polarity during the epithelial-to-mesenchymal transition (EMT) is a hallmark of cancer and metastasis. The key feature of this polarity in epithelial cells is the subdivision of the plasma membrane into apical and basolateral domains, with each orchestrating specific intra- and extracellular functions. Epithelial transport and signaling capacities are thought to be determined largely by the quality, quantity and nanoscale organization of proteins residing in these membrane domains, the apicobasal surfaceomes. Despite its implications for cancer, drug uptake and infection, our current knowledge of how the polarized surfaceome is organized and maintained is limited. Here we used chemoproteomic surfaceome scanning to establish proteotype maps of apicobasal surfaceomes and reveal quantitative distributions of i.a. surface proteases, phosphatases and tetraspanins as potential key regulators of polarized cell functionality. We show further that tumor-suppressor PTEN regulates polarized surfaceome architecture and uncover a potential role in collective cell migration. Our differential surfaceome analysis provides a molecular framework to elucidate polarized protein networks regulating epithelial functions and PTEN-associated cancer progression.

**Summary:** One cell, two functionally different surfaceomes: Chemoproteomic surfaceome scanning reveals quantitative polarization of protein networks across the epithelial cell membrane and unrecognized roles of tumor suppressor PTEN in surfaceome organization associated with cancer progression.

## Introduction

In epithelial tissues, such as the respiratory epithelium and the blood-brain barrier, the plasma membrane has a polarized structure. The resulting apical and basolateral membrane domains act as gateways for signaling molecules and nutrients and are the interfaces for host-pathogen interactions and drug uptake. Remarkably, the functionality and responsiveness of epithelia are specific to each surface. This is exemplified by EGF-mediated signaling, which has long been known to differ upon apical versus basolateral induction (Amsler & Kuwada 1999; Kuwada et al. 1998; Hobert et al. 1999). Presumably, the proteins residing in each membrane domain - here collectively referred to as the apicobasal surfaceome - are decisive for side-specific epithelial operations, and disturbance of their polarized organization leads to altered functioning of the cell. Loss of apicobasal polarity is commonly observed in carcinogenesis and occurs during the epithelial-to-mesenchymal transition (EMT), in which epithelial cells dedifferentiate into the migratory mesenchymal phenotype that promotes metastasis (Thiery et al. 2009). Accordingly, expression aberrations of polarity-controlling proteins are observed in many different cancers (Martín‐Belmonte & Rodríguez‐Fraticelli 2009; Muthuswamy & Xue 2012). Yet, we only have limited knowledge on polarized surfaceome organization and maintenance.

This can be partially attributed to the fact that until recently studies on polarized protein localization and trafficking have been largely based on confocal microscopy and limited to single model proteins (Polishchuk et al. 2004). A few studies have employed mass spectrometry (MS) for protein identification in the polarized membrane, but have focused on the more easily accessible apical surface (Loo et al. 2013; Uchida et al. 2011) or disregarded the quantities of proteins (Yu et al. 2006), a parameter that critically impacts the functional capacity of a membrane. Importantly, proteins do not act on their own in order to fulfill cellular functions but in stable or transient interaction with other proteins (Hartwell et al. 1999). The composition of the proteome and its organization into functional modules through a plethora of protein-protein interactions is defined as the proteotype of a cell (Aebersold & Mann 2016). Knowledge on the proteotype of the polarized surfaceome is key to understanding the differential functioning of apical and basolateral membranes as a basis to resolve mechanisms of pathogenesis (Bausch-Fluck et al. 2019).

Chemoproteomic technology in combination with quantitative MS analysis enables the enrichment and reliable quantification of cell surface proteins (Elschenbroich et al. 2010; Aebersold & Mann 2016; Wollscheid et al. 2009). In this study, we used quantitative chemoproteomic approaches for a detailed characterization of apical and basolateral proteotypes in order to identify key elements of their polarized regulation, and explore how the synergy of local proteins translates into site-specific membrane functionality. In addition, we investigated the perturbation of apicobasal polarity studying the impact of tumor suppressor PTEN on the polarized surfaceome as well as on the epithelial proteotype in the context of carcinogenesis.

## Results

### Chemoproteomic approach for mapping of polarized surfaceomes

In order to map the apicobasal surfaceome quantitatively, we used filter-grown MDCK cells as the best-established *in vitro* system for epithelial polarity (Rodriguez-Boulan et al. 2005) in combination with chemoproteomics and stable isotope labeling of amino acids in cell culture (SILAC) (Ong et al. 2002; Mann 2014). Briefly, 6-day old cultures of filter-grown MDCK cells allowed for specific labeling of apical and basolateral surface proteins with amine-reactive N-hydroxy-succimide (NHS), which we confirmed by confocal microscopy (Fig. S1 B). We employed sulfo-NHS-SS-biotin to specifically label and enrich for apical and basolateral proteins for subsequent MS analysis. To account for the presumably different total protein content of the two membrane domains, we used SILAC cultures to derive apical and basolateral proteins from the same cell growth area and to relatively quantify them by MS-analysis based on the ratio of their heavy and light isotopes (Fig. S1 A). Final normalization for technical variations of heavy and light intensities was accomplished based on the overall heavy-to-light protein ratio measured in the intracellular fraction of each sample. Finally, identified proteins were filtered for evidence for localization at the plasma membrane using UniprotKB and the Surfy surfaceome predictor (Bausch-Fluck et al. 2018) (Fig. S1 C).

### Quantitative protein distribution is the key feature of the apicobasal surfaceome

Using this approach, we were able to confidently quantify about 400 proteins across the polarized membrane of MDCK cells. From these data we generated a quantitative map of the apicobasal surfaceome to characterize key features and functional capacity of the two membrane domains (Fig. 1 A, Table S1, Fig. S2). Interestingly, most of the proteins were found in both surfaceomes. Only a few proteins (around 5%) were exclusively detected apical or basolateral, including several proteins for which the polarized localization has been previously described, as well as a number of novel apical or basolateral markers (Fig. 2 A). Intriguingly, although most surface proteins were generally detected in both membrane domains, more than 60% of them showed a quantitatively polarized distribution (apical:basolateral ratio <40:60 or >60:40). Notably, proteins with a distribution polarized towards the apical surface were substantially less frequent than towards the basolateral surface (average apical:basolateral ratio about 40:60), presumably due to the larger basolateral surface area (Fig. 1 B).

**Fig. 1:**
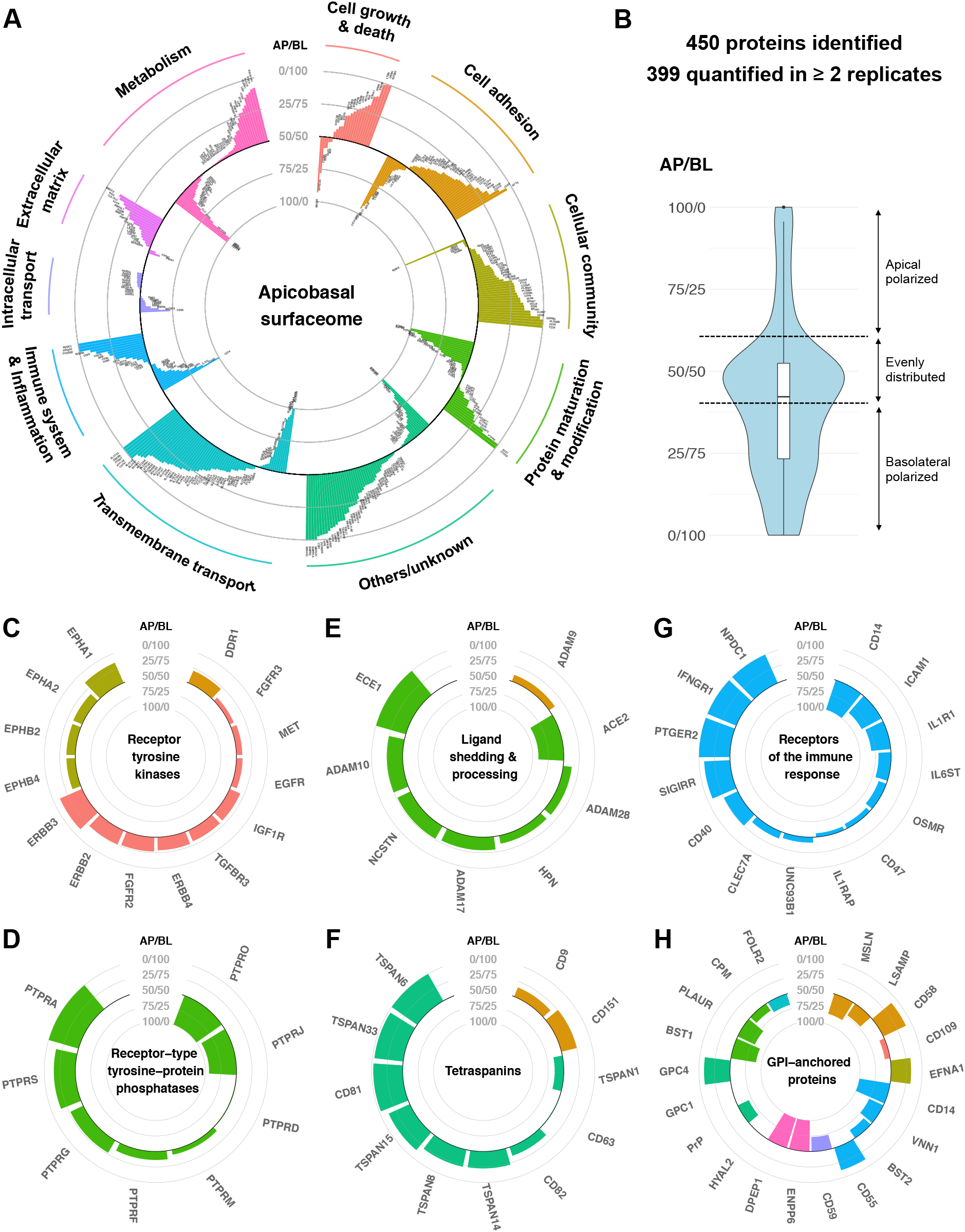
Quantitative apicobasal distribution (AP/BL ratio) of cell surface proteins on filter-grown MDCK cells based on SILAC-complemented chemoproteomics. **(A)** Quantitative map of the apicobasal surfaceome (apical tendencies more central, basolateral tendencies more peripheral in the plot). **(B)** Overall apicobasal protein distribution shows quantitative polarization for more than 60% of the detected proteins with an average ratio of 40:60. **(C-H)** Quantitative maps of different protein classes across the apicobasal surfaceome: (C) receptor-tyrosine kinases, (D) receptor-type tyrosine-protein phosphatases, proteins involved in ligand shedding and processing, (F) tetraspanins, (G) receptors of the immune response, and (H) GPI-anchored proteins. Classification according to the Gene Ontology and KEGG pathway databases (n=4, with 2 SILAC forward and 2 reverse experiments).

**Figure 2:**
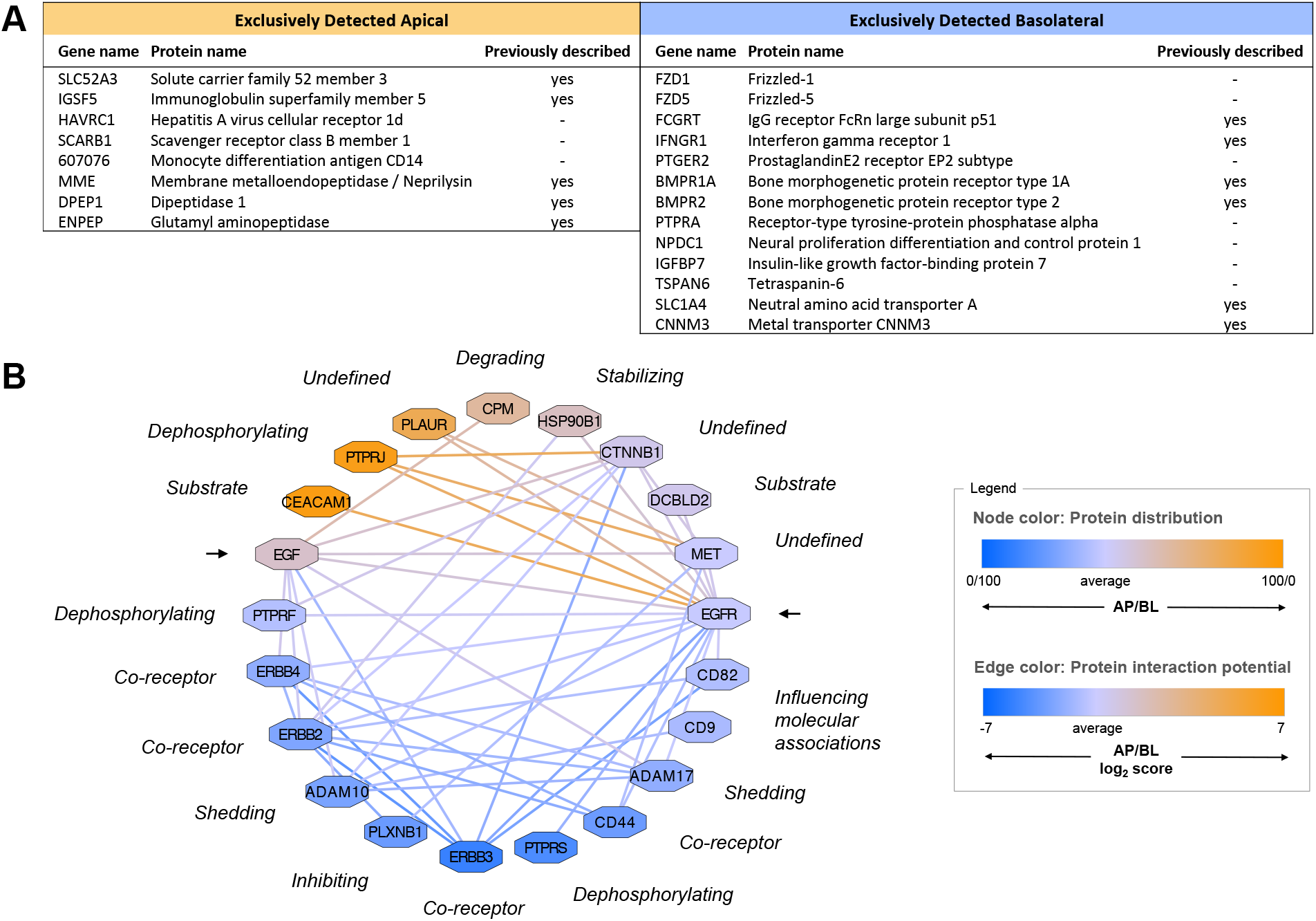
The apicobasal proteotype is determined by polarized protein quantities. **(A)** Only very few proteins were detected exclusively in the apical or basolateral membrane of filter-grown MDCK cells based on SILAC-complemented chemoproteomics (n=4, with 2 SILAC forward and 2 reverse experiments). **(B)** Representation of the polarized interaction network influencing EGF growth signaling, based on quantitative proportions of proteins within the apical versus basolateral membrane. Shown are selected known protein interactions and their functional impact on EGF and EGFR (arrows) according to the STRING and Uniprot databases as well as literature. The quantitative distribution of protein nodes across the apicobasal surfaceome and resulting likelihood of interaction within each membrane domain is color-coded (color scale from orange=apical to blue=basolateral).

We exploited our data to resolve apparent contradictions in the field of apicobasal polarity and trafficking. Protein delivery to each membrane domain requires a polarized intracellular sorting machinery and has been shown to be determined by sorting signals within cargo proteins that are recognized by specific adaptor proteins (Stoops & Caplan 2014; Rodriguez-Boulan & Macara 2014). Despite the proposed function of glycosylphosphatidylinositol (GPI) anchors as apical sorting signals (Lisanti et al. 1989; Rodriguez-Boulan et al. 2005), a previous MS-based study on MDCK cells suggested a non-polarized distribution of the great majority of GPI-anchored proteins (Cortes et al. 2014). The presented quantitative data indicate that although most GPI-anchored proteins are present in both domains, 86% of the detected GPI-anchored proteins do show a polarized distribution ‒ in a quantitative fashion ‒ with a clear tendency toward the apical surface (Fig. 1 H).

### Protein abundances indicate functional capacities of apical versus basolateral membranes

Differences in protein abundance in the apical versus the basolateral membrane are indicative of distinct functions of these membrane domains. For instance, the basolateral surface appears to predominantly mediate cell adhesion and functions involved in cellular community, whereas both apical and basolateral membranes are suggested to fulfill various tasks of the immune and inflammatory system (Figure 1 A). More precisely, we found strongly polarized localizations for different receptors of the immune response, transmembrane transporters and G-protein coupled receptors (Fig. 1 G, Fig. S2). Based on receptor abundance, we can deduce that detection of bacterial lipopolysaccharide via CD14 as well as SCARB1-dependent uptake of lipids and HDL occurs exclusively at the apical cell surface. Further, response to proinflammatory interleukin-1 via the IL1R1 receptor is expected to be mainly mediated through the apical membrane. In contrast, planar polarity (via Wnt signaling) is suggested to be regulated through the basolateral surface, based on localization of frizzled receptors 1 and 5. Also pro-angiogenic signaling of PTGER2 receptor in response to prostaglandin E2 hormone, a pathway currently discussed as a potential drug target for cancer therapy, is only expected from the basolateral side. Conclusively, the quantitative map of the apicobasal surfaceome enables to identify cellular functions that are orchestrated by epithelia in a polarized fashion.

### Polarized distribution of modulatory proteins

Whereas various receptor tyrosine kinases (RTKs) that mediate major cellular functions such as growth signaling (e.g., EGFR and MET) showed rather similar apical and basolateral abundances (Fig. 1 C), many of their modulatory interaction partners were found to be strongly polarized. In particular, about three-fourths of receptor-type tyrosine-protein phosphatases (PTPRs/PTPNs) were detected with substantial polarization, indicating side-specific as well as universal functions of these phosphatases (Fig. 1 D). PTPRs have been shown to dephosphorylate specific sites of different RTKs and thereby have a direct impact on downstream signaling (Meeusen & Janssens 2017; Stanoev et al. 2018). Proteins of the tetraspanin family, which are believed to regulate the function of their interaction partners by membrane compartmentalization (Termini & Gillette 2017), also showed a polarized distribution, mostly with a strong preference for the basolateral domain (Fig. 1 F). Furthermore, our quantification revealed the polarized distribution of several proteolytic enzymes that are required for the shedding of signaling molecules from the cell surface or protein processing in the extracellular space (Fig. 1 E). These findings highlight modulatory proteins that arrange, process, or modify other proteins as key elements of polarized membrane organization.

### Quantitative protein distributions give rise to polarized functional networks

Subcellular compartmentalization is a key regulator of the proteotype, as it generates local networks of specific protein subsets (Thul et al. 2017; Pankow et al. 2019). Accordingly, the quantitative distribution of proteins across the apicobasal surfaceome determines their interaction space within each membrane domain, that may result in differential regulation of apical and basolateral functions. In the context of growth signaling, for instance, we know that apical EGF stimulation induces a different intracellular response than basolateral stimulation (Amsler & Kuwada 1999; Kuwada et al. 1998; Hobert et al. 1999). However, we show here that both, the EGF precursor and its receptor EGFR, are relatively evenly distributed across the polarized membrane (with an apicobasal ratio of 50:50 and 40:60, respectively). Therefore, we created an *in silico* map of the apicobasal protein interaction network exemplified by the EGF/EGFR system (Fig. 2 B) and examined how the differential abundance of known interaction partners in the apical versus basolateral membrane may affect the functioning of this signaling machinery.

Firstly, we found the ADAM10 protease, that is known to cleave the EGF precursor from the cell surface to release the active growth factor, mainly in the basolateral domain. In contrast, the CPM peptidase, which is believed to control growth factor activity by hydrolysis, predominantly localizes to the apical surface. This suggests that EGF of epithelial origin exhibits mostly basolateral activity under physiological conditions, despite the unpolarized distribution of the precursor.

Furthermore, we found a number of RTK co-receptors and phosphatases, previously shown to modulate phosphorylation and function of the EGF receptor, to be quantitatively polarized. Unlike EGFR (ErbB1), other members of the ErbB growth factor receptor family (ErbB2-4) were detected as differentially abundant in the apical versus the basolateral membrane domain. ErbB receptors are known to form homo- and heterodimers upon binding of particular ligands and induce alternative cross-phosphorylation and intracellular signaling pathways depending on the dimer formed (Yarden & Sliwkowski 2001). In addition, several PTPR phosphatases that are known to interact with EGFR showed a polarized distribution. Based on protein abundance, the formation of EGFR/EGFR homodimers and the regulation by PTPRJ are more likely to occur in the apical domain, whereas EGFR/ErbB3 heterodimers and interaction with PTPRS and PTPRG may have higher prevalence in the basolateral domain. This distinct interaction potential with other RTKs and phosphatases suggests that differentially phosphorylated proteoforms of EGFR are generated upon apical versus basolateral EGF binding to modulate downstream signaling.

Additionally, apicobasal EGF signaling may be influenced by crosstalk with other receptor signaling pathways and tetraspanins such as CD9 and CD82, which are thought to determine the substrate specificity of ADAM proteases. Conclusively, cellular responsiveness to e.g. growth factors may be tailored to the specific functions of the apical and basolateral surface by distinct interaction networks that are dominated by polarized distributions of modulatory proteins.

### Tracking of polarized protein trafficking suggests intertwining of sorting routes

The current knowledge on mechanisms of polarized protein sorting does not reflect the diverse quantitative protein distributions we observed. Therefore, we explored the applicability of chemoproteomic approaches to directly track polarized protein delivery. We employed pulsed SILAC in combination with a membrane-fixation strategy based on tannic acid. Tannic acid has been shown previously to block the fusion of intracellular vesicles with the plasma membrane and to fix the apical or basolateral membrane of filter-grown MDCK cells in a side-specific manner (Polishchuk et al. 2004). We confirmed full integrity of treated cell layers by confocal microscopy (Fig. 3 A). After tannic acid treatment to block vesicular delivery to one membrane domain, we switched the amino acid source to heavy isotopes, allowing us to track trafficking of newly synthesized (heavy) proteins to the other membrane domain (Fig. 3 B). Four hours after blocking delivery to the basolateral membrane, we observed that the heavy version of most proteins accumulated in the apical membrane compared to the control that was not treated with tannic acid (Fig. 3a). In contrast, blocking apical protein delivery had no significant effect on protein delivery to the basolateral surface, except for the transcytotic LDL receptor (Fig. 3 C). Hence, most proteins in the biosynthetic pathway were redirected to the apical surface if basolateral delivery was denied but not the other way around. Interestingly, we also found that turnover of a few surface proteins was location-specific with more rapid incorporation of the heavy version in one membrane domain compared to the other (Fig. S5). In conclusion, this experiment demonstrated that chemoproteomic approaches are also potent tools to dissect polarized protein trafficking and showed an intertwining of apical pathways with basolateral routes.

**Fig. 3:**
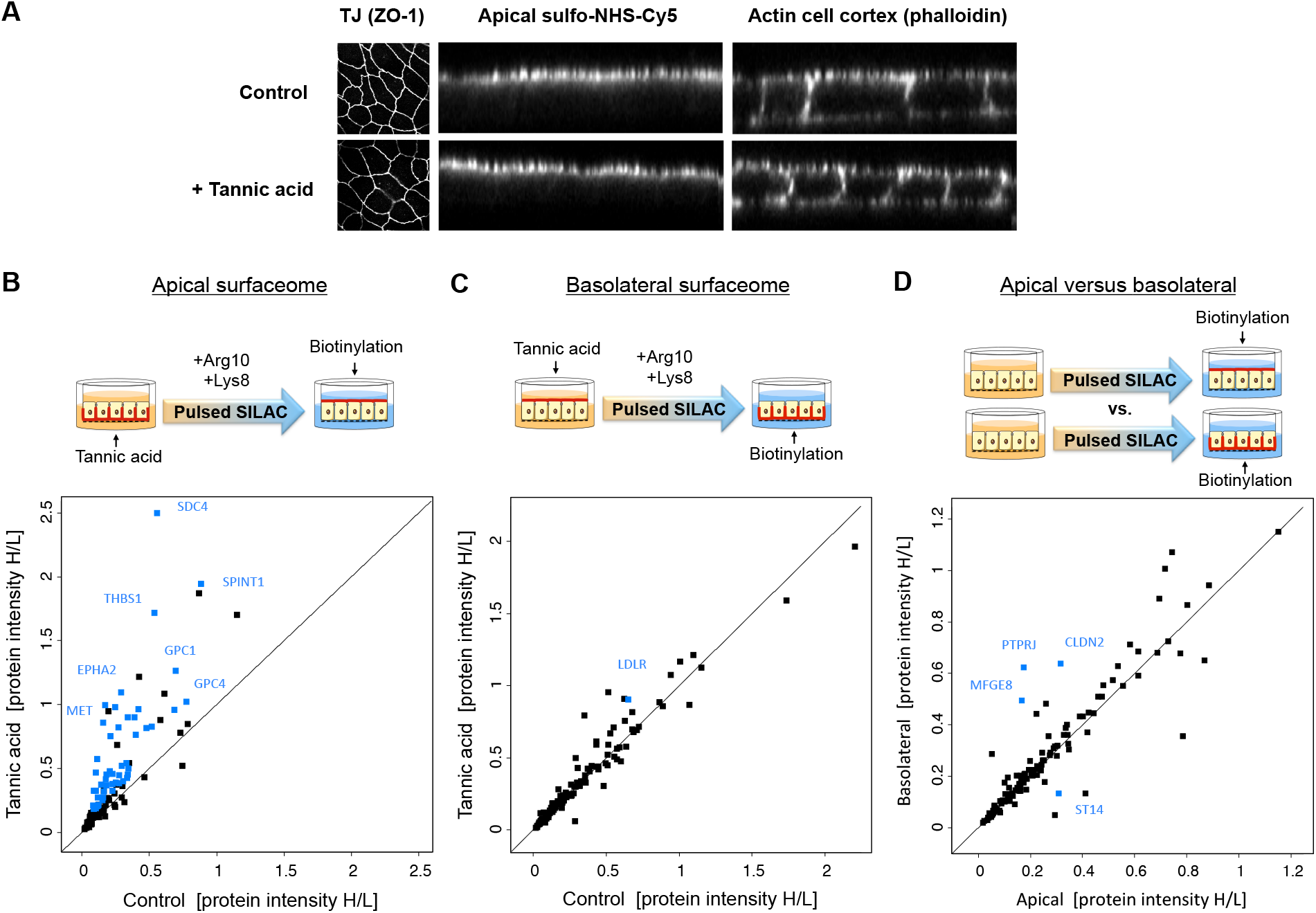
Pulsed SILAC experiment reveals flexibility in the delivery of newly synthesized proteins to the apical and basolateral surfaces upon side-specific blocking by membrane fixation using tannic acid. **(A)** Confocal microscopy imaging of filter-grown MDCK cells shows intact cell layer integrity 4 h after tannic acid treatment (0.5% for 10 min) compared to the untreated control, with no effect on the expression of the tight junction (TJ) protein ZO-1 (left panel) nor on the side-specific cell surface labeling with an NHS-conjugate (center panel). Staining of the actin cell cortex by phalloidin was used as a reference for cellular localization (right panel). **(B)** Delivery of many newly synthesized (heavy) proteins was increased to the apical surface upon basolateral tannic acid treatment. **(C)** Almost no changes in protein delivery to the basolateral surface proteins were observed upon apical tannic acid treatment. **(D)** Comparison of the heavy-to-light ratio of apical and basolateral surface proteins indicates that some newly synthesized proteins integrate more rapidly into either the apical or basolateral membrane domain, suggesting side-specific turnover rates for these proteins. Blue dots mark proteins that show statistically significant differences between conditions (shown are median values, n=3, t-test FDR=0.05, s0=0.1).

### Impaired PTEN function causes alterations in the polarized surfaceome

After establishing the quantitative composition of the apicobasal surfaceome in a physiological setting, we aimed to understand polarity-associated defects that may contribute to cancer progression. PTEN is one of the most frequently mutated tumor suppressors in humans. Besides its established role as antagonist of PI3-kinase-dependent growth signaling, PTEN has also been shown to play a role in EMT (Hu et al. 2019) and to be a determinant of epithelial polarity. PTEN functions as a protein and lipid phosphatase known to e.g. dephosphorylate phosphatidylinositol (3,4,5)-trisphosphate (PtdIns(3,4,5)P_3_) to phosphatidylinositol (4,5)-biphosphate (PtdIns(4,5)P_2_). It has been shown that PtdIns(3,4,5)P_3_ plays a role in establishing the basolateral domain and is excluded from the apical membrane, whereas PtdIns(4,5)P_2_ as well as PTEN localize to the apical membrane in MDCK cyst cultures. This localization is required for the recruitment of the apical polarity complex, which coordinates the formation of tight junctions essential for polarity (Gassama-Diagne et al. 2006; Martin-Belmonte et al. 2007). We hypothesized that PTEN function is also crucial for the polarized localization of cell surface proteins.

In order to investigate the effect of the phosphatase activity of PTEN on apicobasal surfaceome composition, we treated filter-grown MDCK cells with the small-molecule inhibitor SF1670 in low concentration (1 μM versus IC50 = 2 μM to retain target specificity). Using short incubations (4 h), we initially targeted the acute response of polarized protein trafficking, while avoiding cellular contra-regulation on the level of protein expression. Importantly, the treatment had no impact on neither the integrity of the cell layer (Fig. 4 A) nor the cellular abundance of most proteins under these conditions (Fig. 5 A). In a label-free MS experiment we found that the quantities of a number of apical surface proteins were altered upon PTEN inhibition compared to the mock-treated control, albeit most proteins remained unchanged. Among proteins with increased abundance were receptors that regulate cellular growth and cohesion, whereas a few proteins involved in apicobasal polarity and transport to the cell surface were downregulated (Fig. 4 B). This effect was rescued by simultaneous inhibition of the PI3-kinase (Fig. 4 C), indicating a PtdIns phosphorylation-dependent trafficking mechanism of altered apical surface proteins.

**Fig. 4:**
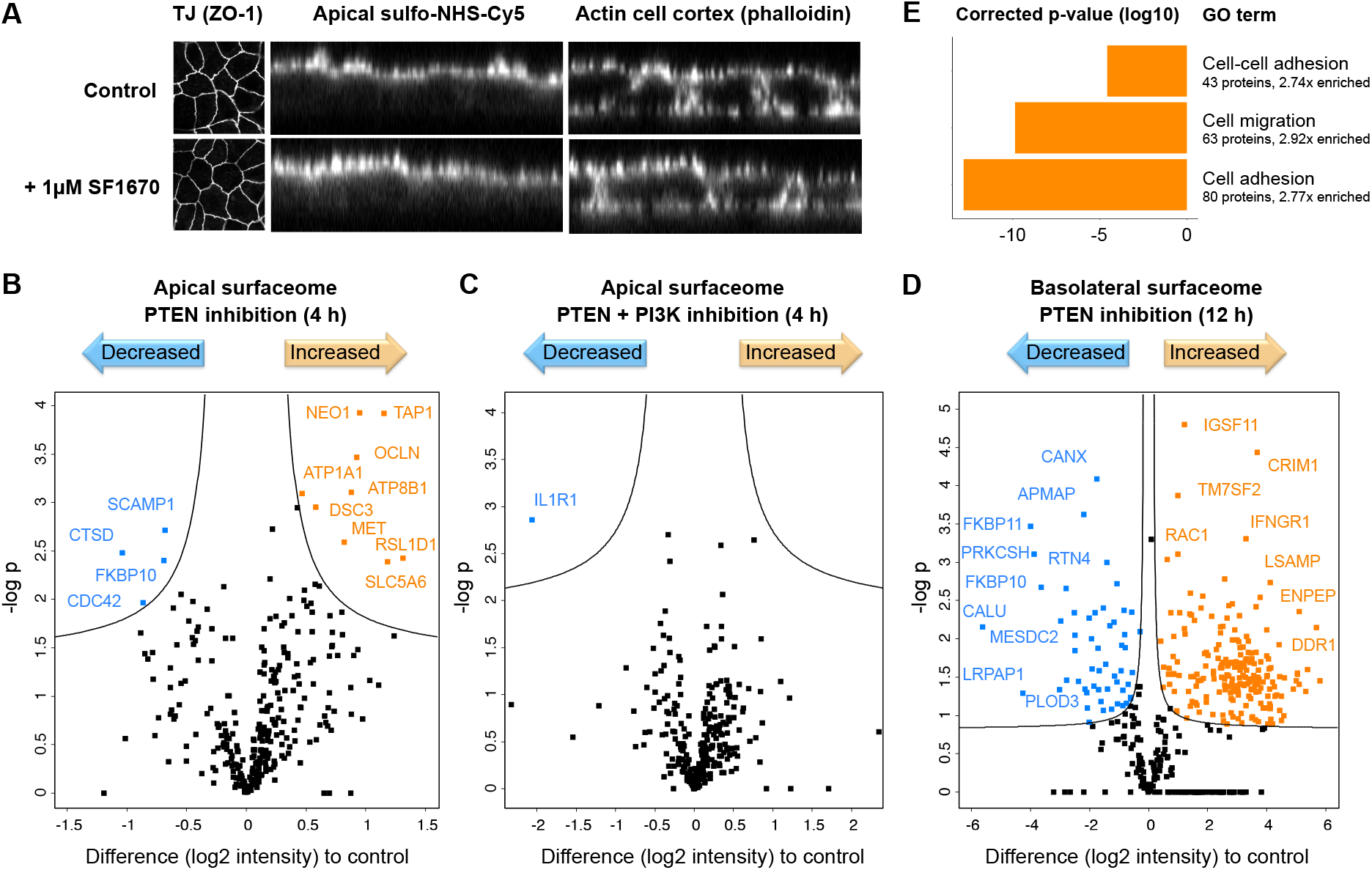
Impaired PTEN function impacts the polarized surfaceome. **(A)** Representative confocal microscopy images of filter-grown MDCK cells showed intact cell layer integrity upon PTEN inhibition by 1 μM SF1670, with no effect on the expression of the tight junction (TJ) protein ZO-1 (left panel) nor on the side-specific cell surface labeling with an NHS-conjugate (center panel). Staining of the actin cell cortex by phalloidin was used as a reference for cellular localization (right panel). **(B)** Label-free quantification of apical surface proteins upon PTEN inhibition by 1 μM SF1670 for 4 h revealed alterations in protein abundance. **(C)** Simultaneous PTEN and PI3-kinase-inhibition by 1 μM SF1670 and 1 μM LY294002, respectively, diminished the effect. **(D)** PTEN inhibition caused substantial quantitative changes in the basolateral surfaceome composition. Proteins with significantly altered abundance compared to the control are marked in orange (increased) and blue (decreased) (n=3, t-test FDR=0.05, s0=0.1). **(E)** Gene ontology analysis of basolateral proteins that showed increased levels upon 12h PTEN inhibition revealed enrichment of proteins involved in cell adhesion, cell-cell adhesion and cell migration (Fisher’s exact test with Bonferroni correction).

**Fig. 5:**
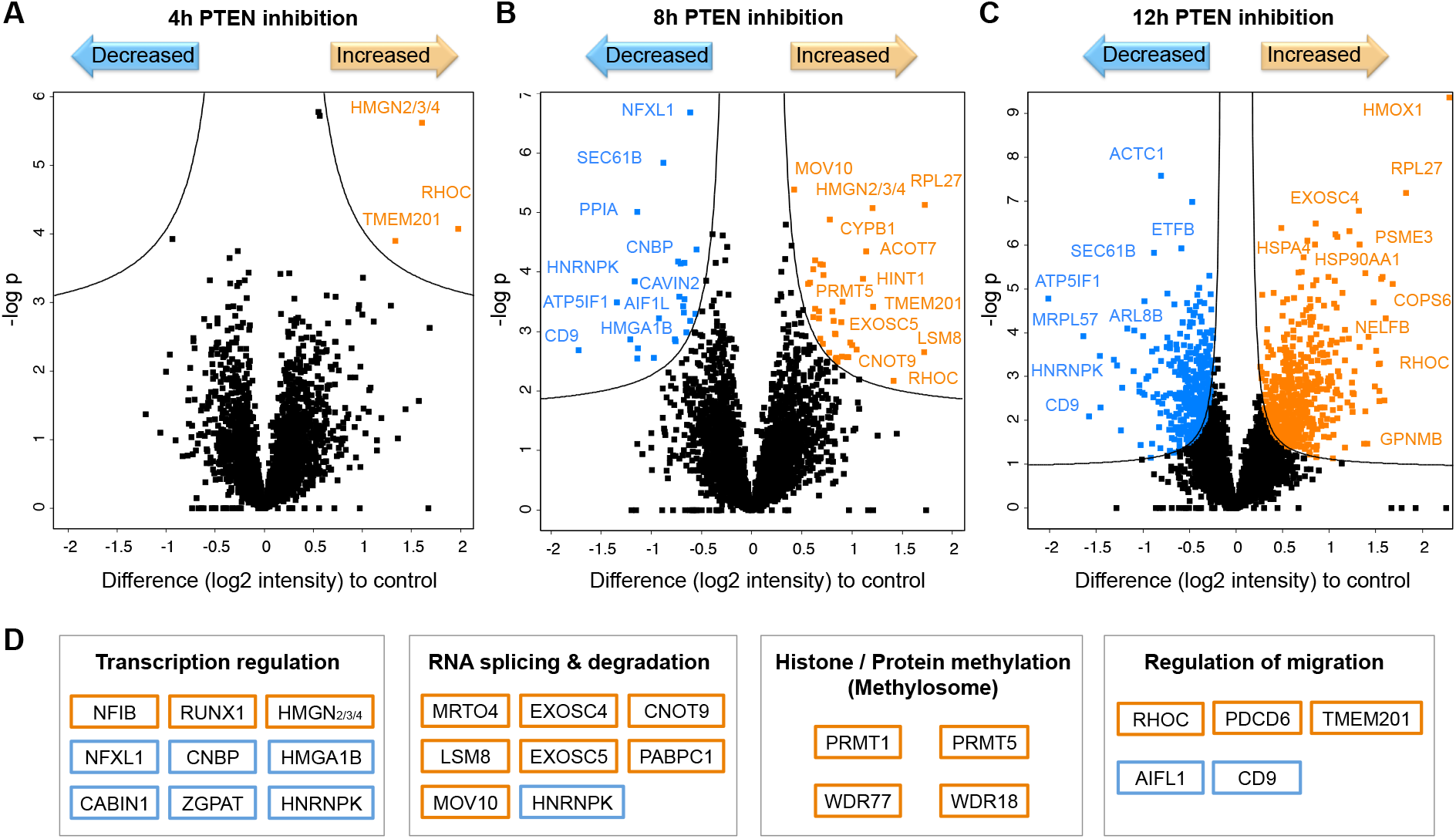
Protein expression changes upon PTEN inhibition in polarized MDCK cells indicate a massive cellular reorganization. **(A-C)** Label-free quantification of the cellular proteotype over a time course of PTEN inhibition by 1 μM SF1670 for 4 h, 8 h and 12 h. **(D)** Functional classes of selected proteins that were found with altered abundance after 8 h of PTEN inhibition as potential drivers of proteotype reorganization. Proteins with significantly altered abundance compared to the control are marked in orange (increased) and blue (decreased) (n=6, t-test FDR=0.05, s0=0.1).

In line with apical localization of PTEN, the basolateral surfaceome was unaffected by PTEN inhibition after 4 h (data not shown). However, at a later stage of PTEN inhibition (12 h) we observed substantial changes in basolateral surfaceome composition (Fig. 4 D). For instance, we detected enhanced surface (but not overall) expression of growth factor receptors such as EGFR, MET, IGF1R and ERBBs. Intriguingly, Gene Ontology analysis of proteins with elevated abundance revealed an enrichment of proteins that mediate cell adhesion and migration, suggesting that cells gained motility (Fig. 4 E). At the same time, increased expression of cell-cell adhesion proteins, including epithelial markers such as CDH1/E-cadherin, indicated enhanced cell cohesion. Among downregulated proteins were for instance regulators of extracellular matrix organization. Strikingly, a number of proteins that are known to play important roles in cancer progression and invasion such as Podocalyxin and inhibitory Calumenin were found upon PTEN inhibition with increased or decreased basolateral levels, respectively. Taken together these data suggest that tumor-suppressor PTEN does not only regulate apical surfaceome maintenance but ultimately also cellular motility through modulation of the basolateral surfaceome.

### Impaired PTEN function causes massive proteotype reorganization

In order to resolve the underlying mechanisms of surfaceome changes following loss of PTEN functionality, we performed proteotype analyses of polarized MDCK cells in a time course of inhibitor treatment (Fig. 5). In a label-free MS experiment we identified about 4260 proteins in total. After 4 h of inhibitor treatment we only observed abundance changes of a few proteins (Fig. 5 A), whereas at a later stage (8 h) the inhibitor treatment led to quantitative alterations of more than 60 proteins (Fig. 5 B). After 12 h of PTEN inhibition profound expression changes, comprising roughly one-fifth of the detected proteins, indicated a massive reorganization of the cellular proteotype (Fig. 5 C).

The earliest regulated proteins, that also persisted throughout the time course and are thus potential initiators of the observed effects, were RHOC, TMEM201 and a HMGN protein (2, 3 or 4). RHOC and TMEM201 are known to regulate cell junction assembly and cell migration (Gene Ontology), in line with subsequent basolateral expression of respective proteins described above. HMGN proteins are binding to nucleosomes for transcriptional regulation, with the downstream effects being unclear.

At the intermediate time point of 8 h we identified several regulators of gene transcription and RNA processing/degradation as well as known regulators of migration, that might be driving proteotype reorganization (Fig. 5 D). Additionally we found components of the methylosome complex among upregulated proteins, which mediates arginine methylation of histones and other protein targets. Both methyltransferases, PRMT1 and PRMT5, have been shown to be overexpressed in many cancers and to regulate cancer cell migration and invasion (Chen et al. 2017; Avasarala et al. 2015; Yang & Bedford 2013). Transcriptional regulation by histone methylation may therefore play a crucial role for subsequent effects of PTEN inhibition.

Among proteins with elevated levels after 12 h of inhibitor treatment were proteins involved in RNA processing and protein translation, as well as chaperones (e.g., HSP90) and proteins associated with extracellular exosomes. Importantly, we did not detect known markers of EMT or mesenchymal differentiation at any point, in agreement with sustained expression of cell-cell contacts. However, we found a number of established drivers of cancer progression and invasion (e.g., mTOR, STAT1/5, RHOC, ARF6, HMGB, CRK) with increased expression levels as well as known suppressors (e.g. AIFL1, SLIT3) with decreased expression levels upon PTEN inhibition.

Taken together, our data show that impaired PTEN function does not lead to loss of epithelial cell polarity but to a global reorganization of the polarized surfaceome as well as the intracellular proteotype, that resembles a migratory but not mesenchymal phenotype.

## Discussion

In this study we used chemoproteomic approaches to map the apicobasal surfaceome in an *in vitro* model and to investigate perturbations associated with carcinogenesis. We specifically identified quantitative differences in protein composition as a major factor for epithelial surface organisation. This was particularly the case for modulatory proteins that organize, process, or modify other proteins in order to regulate their activity. A recent study used a similar approach to quantify apicobasal protein distributions in MDCK cells for research on polarized protein sorting (Caceres et al. 2019). The data are remarkably consistent with our map of the apicobasal surfaceome, confirming our results by mass spectrometry as well as immunoblotting. The two studies establish quantitative chemoproteomics as a powerful tool to investigate epithelial polarity.

Differential protein abundances suggest that there are distinctive interaction potentials for each protein within the apical versus the basolateral domain. This is likely to influence the nanoscale organization of proteins, perhaps with modulatory proteins and co-receptors, which is believed to be crucial for physiological cell signaling (Bausch-Fluck et al. 2019). Differential apicobasal EGF signaling has been associated with domain-specific EGFR binding of intracellular signaling proteins (Amsler & Kuwada 1999). We hypothesize that the upstream molecular basis for such observations is a polarized receptor nano-scale organization resulting in the generation of side-specific (phospho) proteoforms that recruit different downstream signaling proteins. Future studies may follow up on the emergence of specific apical and basolateral phosphoforms during signal transduction, and verify domain-specific interactions with regulatory proteins by proximity-labeling or other strategies (Li et al. 2014). It will also be interesting to investigate how for instance distinct glycoforms or additional mechanisms of membrane compartmentalization (e.g., cilia) may contribute to apical and basolateral functionality. Moreover, the surfaceome map could be extended by the incorporation of additional types of data (e.g., RNA-seq data from polarized cells to identify splice variants).

Our data on quantitative surfaceome polarity emphasize the complexity of polarized protein trafficking. The occurrence of known apical and basolateral sorting signals is usually not limited to one membrane, and it has been shown previously that apicobasal sorting of a particular protein can depend on multiple sorting signals (Alonso et al. 1997; Youker et al. 2013). In this study we showed that most proteins can be redirected to the apical surface if basolateral delivery is inhibited, indicating an intertwining of apicobasal trafficking routes. Therefore, we propose a model in which apical sorting machineries act competitively with the basolateral (default) route in order to establish a specific protein distribution. Competitive sorting might be a beneficial mechanism to enable the “flexible phenotype” of epithelia in distinct tissues and physiological conditions. In-depth bioinformatic analyses of the data generated in this work may identify patterns in apical versus basolateral protein features required for such a sorting system.

Our findings concerning tumor suppressor PTEN demonstrate that PTEN is implicated in apical localization of cell surface proteins, in line with previous studies of PTEN functions in polarity (Gassama-Diagne et al. 2006; Martin-Belmonte et al. 2007). Further we provide evidence that inhibited PTEN activity causes increased expression of known drivers of cancer cell migration and invasion, followed by a substantial reorganization of the epithelial proteotype as well as the basolateral surfaceome that suggest cell motility. While these findings draw parallels to EMT, we did not find any evidence for a loss of cell cohesion and polarity, or alterations of major EMT markers. This phenotype may resemble collective cell migration, describing the movement of cell sheets rather than single cells, which has been shown to play a role in wound healing as well as tumor metastasis (Mayor & Etienne-Manneville 2016). PTEN has been shown previously to play a role in leukocyte motility and in restricting cell migration in wound healing (Lacalle et al. 2004; Cao et al. 2011; Squarize et al. 2010). In fact, E-cadherin - as a typical marker lost during EMT that we found to be upregulated upon PTEN inhibition - has been shown to be essential for collective cell invasion (Labernadie et al. 2017). Interestingly, higher concentrations of PTEN inhibitor led to cell detachment in large patches (personal observation). Collectively, our findings point towards an acquired motility, potentially collective cell migration, upon PTEN inhibition and identify several potential driver proteins hereof. Future studies may validate these findings by PTEN knock-down experiments and follow up on them by detailed phenotyping of PTEN-impaired epithelia, including e.g. migration assays, exosome analysis and profiling of histone methylation.

Overall, the map of the apicobasal surfaceome generated here enabled system-wide insights into the protein organization and maintenance required for the polarized operation of epithelial cells. Providing concepts of epithelial plasticity and the role of tumor suppressor PTEN, our findings will facilitate future studies of apical and basolateral functions and mechanisms of cancer progression.

## Acknowledgments

We are grateful to Maria Pavlou for her feedback at all stages of the project as well as on the manuscript. We thank Carl Philipp Zinner for his support in implementing data visualisation. We acknowledge Jacqueline Wyatt for editing the manuscript, and the Hugo Stocker / Ernst Hafen lab for access to microscopes. We thank Jason Mercer, Yohei Yamauchi and Ruedi Aebersold for scientific discussions and for providing cell lines.

We acknowledge the Swiss National Science Foundation (grant 31003A_160259; for B.W.) for their generous support enabling this project.

The authors declare no competing financial interests.

## Author Contributions

A.K. and B.W. conceived the study and wrote the manuscript. A.K. designed and conducted laboratory experiments, and performed data acquisition and analysis.

## Materials & methods

### Chemicals

All chemicals and cell culture reagents were from Sigma-Aldrich unless stated otherwise.

### Cell lines

MDCK II cells were a gift from Yohei Yamauchi (University of Bristol).

### Mammalian cell culture, SILAC labeling, and polarized culture

Cells were grown at 37 °C and 5% ambient CO_2_ to approximately 80% confluence in 140 × 20 mm dishes (Nunclon^TM^Delta Airvent, Thermo Fisher) with Dulbecco’s Modified Eagle’s Medium, high glucose, GlutaMAX^TM^supplement (DMEM, Thermo Scientific), 10% fetal bovine serum (FBS; BioConcept), and 1% penicillin-streptomycin. For SILAC labeling, cells were grown in SILAC DMEM high glucose with stable glutamine and without arginine and lysine (Pan Biotech), supplemented with 15% dialysed FBS (Pan Biotech), 1% penicillin-streptomycin, 10 mM HEPES (Gibco), non-essential amino acids (Gibco), 200 mg/L L-proline, 42 mg/L L-arginine or L-arginine[10] and 73 mg/L L-lysine or L-lysine[8] (Silantes) for 7 days, and the incorporation rate of heavy amino acids was determined to be 95-98% by MS analysis. For polarized culture, 5×10^6^ or 0.5×10^6^cells, respectively, were seeded into 75-mm or 12-mm polycarbonate Transwells^TM^with 0.4 μm pore size (Corning) and cultured for 6-7 days with medium change every second day.

### Confocal microscopy imaging

To evaluate the tightness of filter-grown cell cultures, cells were labeled at the apical or basolateral surface using 1 mM sulfo-NHS-Cy5 (Lumiprobe) in PBS, pH 8.0 for 15 min on ice in the dark and washed three times with PBS before fixation. For all microscopy samples, cells were fixed in 4% paraformaldehyde for 10 min at room temperature and permeabilized with 0.1% TritonX-100 in blocking buffer (1% FBS, 1% BSA in PBS with 0.02% sodium azide) for 10min at room temperature. Samples were blocked for 1 h at room temperature or overnight at 4 °C in blocking buffer. Cells were subsequently incubated with mouse anti-ZO1-AF555 antibody (Thermo Invitrogen MA3-39100-A555, 1:100) for 1 h with shaking. Nuclei were stained with 1 μg/ml Hoechst (Molecular probes H1399), and F-actin was stained with phalloidin-iFluor488 (Abcam, 1:1000) for 15 min at room temperature. Samples were fixed with 4% paraformaldehyde for 10 min at room temperature and mounted with Prolong Gold Antifade reagent mounting medium (Molecular Probes). Images were taken with a Leica TCS SP2 confocal microscope and processed using the FIJI software.

### Tannic acid membrane fixation and pulsed SILAC

Filter-grown SILAC-light cells were washed once with pre-warmed PBS and incubated with 0.5% tannic acid in pre-warmed PBS in the upper or lower chamber of the Transwell for 10 min at room temperature. Cells were then washed with PBS and incubated for 4 h with SILAC-heavy medium at 37 °C.

### Inhibitor treatments

Filter-grown cells were treated with 1 μM SF1670 (Lucerna Chem) and/or 1 μM LY29400 in 20 ml serum-free DMEM, 1% penicillin-streptomycin per Transwell (8 ml apical; 12 ml basal) at 37 °C for the indicated incubation times, and inhibitor solution was refreshed every 4 hours (0.002% DMSO end concentration). In a control, cells were mock-treated with 0.002% DMSO.

### Cell surface protein enrichment

For side-specific enrichment of apical and basolateral proteins, filter-grown MDCK cells were labeled with a sulfo-NHS-SS-biotin conjugate (Pierce™ Premium Grade, Thermo Scientific). For this purpose, cultures were placed on ice and washed twice with ice-cold PBS. Cells were rinsed on the side to be labeled with cold PBS, pH 8.0, and the opposite chamber was filled with 25 mM Tris, pH 8.0. The apical or basolateral cell surface was then labeled with 1 mM sulfo-NHS-SS-biotin for 15 min on ice. Cells were rinsed on the labeled side with 25 mM Tris, washed with PBS, and harvested by scraping in PBS and centrifugation (5 min at 300 rpm and 4 °C).

Cells were lysed on ice in lysis buffer (50 mM AmBic, 0.25% RapiGest, cOmplete protease inhibitor cocktail (Roche)) by sonication (Dr. Hielscher sonicator, 30 sec at 80% amplitude and 80% cycle time). Biotinylated proteins were enriched on 100 μl streptavidin Ultralink beads (Thermo Scientific) for 1.5 h at room temperature with rotation. Beads were washed 5 times each with 1 ml of 5 M NaCl, StimLys buffer, 80% isopropanol, 100 mM sodium bicarbonate, pH 11, and 50 mM AmBic using Bio-Spin® columns (BioRad) in combination with a vacuum manifold. Washed beads were transferred to 0.5-ml Eppendorf tubes, and proteins were eluted in 250 μl 50mM AmBic, 5 mM TCEP and 0.1% Rapigest by rotation for 1 h at 37 °C. Eluate was collected and combined with an additional bead wash performed with 250 μl 50 mM AmBic.

### Sample preparation for MS analysis

Protein samples were reduced with 5 mM TCEP for 30 min at room temperature, alkylated with 5 mM iodoacetamide for 30 min in the dark and digested overnight with trypsin (Promega; 1 μg for cell-surface-enrichment samples, 1:100 for lysate samples) at 37 °C, 300 rpm. Samples were acidified to pH 2 by addition of formic acid (FA) and centrifuged (10 min at 16,000 rpm at room temperature). Peptides were desalted on a reverse-phase C18 column (Nest Group) and eluted with 50% acetonitrile (ACN), 0.1% FA. Solvent was evaporated using a SpeedVac® Concentrator (Thermo Scientific). Peptides were resuspended in 5% ACN, 0.1% FA, and iRT retention time peptides (Biognosys) were spiked in for subsequent LC-MS/MS analysis.

### LC-MS/MS analysis

Peptides were analysed on a Orbitrap Fusion mass spectrometer (Thermo Scientific) equipped with a nano-electrospray ion source (Thermo Scientific) and coupled to a nano-flow high pressure liquid chromatography (HPLC) pump with an autosampler (EASY-nLC II, Proxeon). Peptides were separated on a reversed-phase chromatography column (75-μm inner diameter PicoTip™ Emitter, New Objective) that was packed in-house with a C18 stationary phase (Reprosil Gold 120 C18 1.9 μm, Dr. Maisch). Peptides were loaded onto the column with 100% buffer A (99.9% H_2_O, 0.1% FA) at 800 bar and eluted at a constant flow rate of 300 nl/min with a gradient of buffer B (99.9% ACN, 0.1% FA) with a subsequent wash step at 90% buffer B. For the analysis of cell-surface protein enrichment samples and the corresponding unbound fractions, 1 μg of peptides were separated on a 15-cm column with a 90-min linear gradient of 5-35% B, followed by a 5-min gradient to 50% B and a 4-min gradient to 90% B. For the analysis of lysates, 3 μg of peptides were separated on a 45-cm column with a 200-min linear gradient of 5-35% B, followed by a 18-min gradient to 50% B and a 10-min gradient to 90% B. Between batches of runs, the column was cleaned with two steep consecutive gradients of ACN (10%-98%).

The MS was operated in data-dependent acquisition mode with a cycle time of 3 s and using all parallelizable time for ion injection. High-resolution MS scans were acquired in the Orbitrap (120,000 resolution, automatic gain control target value 2×10^5^) within a mass range of 395 to 1500 m/z. Precursor ions were isolated in the quadrupole with an isolation window of 2 m/z and fragmented using higher-energy collisional dissociation to acquire MS/MS scans in the Orbitrap (30,000 resolution, intensity threshold 2.5×10^4^, target value 2×10^5^). Dynamic exclusion was set to 30 s. Instrument performance was evaluated on the basis of regular quality control measurements using a yeast lysate and the iRT retention time peptide kit (Biognosys).

### Data analysis and visualisation

MS data were analysed using the MaxQuant software (version 1.6.0.16) using default settings if not stated otherwise. Searches were performed against the canine UniProt FASTA database (June 2018) and a contaminant database with a false discovery rate (FDR) of 1% at protein and peptide level. For the analysis of cell-surface samples, the NHS modification (C_5_O_2_SNH_7_=145.0197Da at K) was selected as additional variable modification, which was also used for quantification, and three missed cleavages were allowed. For SILAC-based quantification, multiplicity was set to 2, with Arg10 and Lys8 as heavy labels. For label-free quantification (LFQ) of proteome samples, “MaxQuant LFQ” was used. The “match between runs” function was enabled in all analyses.

Bioinformatics analysis was performed with Perseus (version 1.6.0.7) and Microsoft Excel. In order to filter out probable intracellular contaminants in the cell-surface-enriched samples, a “canine surfaceome” list was established, containing all canine proteins annotated as “Cell membrane”, “GPI-anchor”, and “secreted” in UniProtKB as well as the predicted surface proteins contained in the bronze set of “Surfy” (Bausch-Fluck et al. 2018), translated from human to canine IDs (Fig. S1 C). For apicobasal quantification, apical and basolateral fractions of detected proteins were calculated from heavy and light intensities, normalized to the overall heavy-to-light ratio measured in unbound fractions of the protein enrichment. Data were filtered for quantification in at least two out of four replicates and are presented as medians of all replicates. As a measure of the variation of the quantification across replicates, the median absolute deviation (MAD) was calculated. Proteins with a median value of 0 (only basolateral) or 1 (only apical) were only considered as exclusively apical and basolateral proteins, respectively, if MAD<10%. For label-free surfaceome analyses, unfiltered MS intensities were median-normalized and filtered for at least 2 unique peptides and 5 MS/MS counts. Statistical significance of results was determined by two-sided t-test (FDR=0.05, s0=0.1). Functional annotations of proteins were made manually, based on information from UniProtKB and Gene Ontology. Gene ontology enrichment analyses of biological processes were conducted against the surfaceome list as a reference and using Fisher’s exact test with Bonferroni correction. Known protein-protein interactions were extracted from the STRING database using a minimum score of 0.4 excluding text mining. A relative apicobasal interaction potential score was calculated based on the quantitative protein distribution using the following formula: log_2_ [(AP_Node1_ × AP_Node2_) / (BL_Node1_ × BL_Node2_)], with AP being the apical fraction and BL being the basolateral fraction of the total surface pool of a protein. Data were visualized using the Perseus platform, Cytoscape, BioVenn and R.

**Supplementary material.** Fig. S1 shows details of the chemoproteomic approach used to map polarized surfaceomes; schematic of the experimental workflow; confocal images of side-specific labeling; assembly of in silico surfaceome list for data filtering; overlap of identified proteins across replicates; variation of quantification across replicates. Fig. S2 shows the quantitative apicobasal distribution of selected protein classes. Table S1 contains a list of all proteins quantified across the apicobasal surfaceome.

