## Supplementary materials for "Apicobasal surfaceome architecture encodes for polarized epithelial functionality and depends on tumor suppressor PTEN"

### Supplementary material

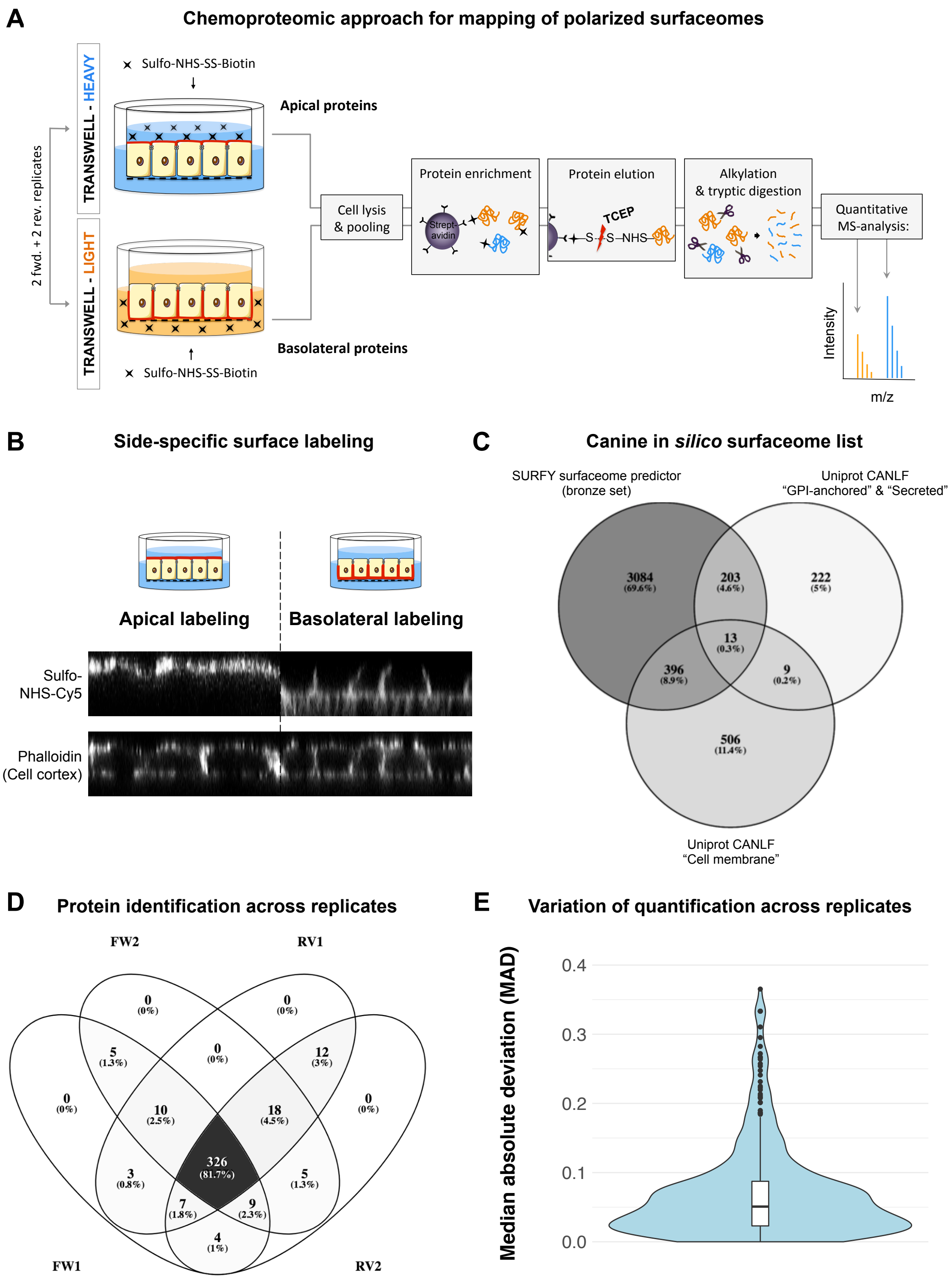

**Figure S1:** Chemoproteomic approach to map polarized protein distributions across the apicobasal surfaceome. **(A)** Workflow: MDCK cells are metabolically labeled with light and heavy isotopes of amino acids using SILAC and polarized in Transwell™ culture. Apical and basolateral surface proteins are tagged in differentially labeled SILAC cultures using a sulfo-NHS-SS-biotin conjugate. Heavy and light cell lysates are mixed, and biotinylated surface proteins are enriched on streptavidin beads. Enriched proteins are eluted by reduction of the disulfide-bond of the biotin conjugate, alkylated and digested with trypsin, and quantified by mass spectrometry. Proteins derived from the apical and basolateral surface are quantitatively compared based on their heavy-to-light ratio. Experiments were conducted in quadruplicates, including two forward and two reverse SILAC replicates. **(B)** Side-specific protein labeling with sulfo-NHS-Cy5 on the apical and basolateral surfaces of filter-grown MDCK cells was confirmed by confocal microscopy imaging. Phalloidin staining of F-actin was used to visualize the cell cortex as a reference. **(C)** Assembly of a canine *in silico* surfaceome list for data filtering using UniprotKB entries annotated as “Cell membrane”, “GPI-anchor”, and “secreted”, as well as the predicted surface proteins contained in the bronze set of “Surfy” (Bausch-Fluck et al. 2018). **(D)** Overlap of identified proteins across replicates and **(E)** a low median absolute deviation (with a median of 5%) shows high reproducibility of the apicobasal protein quantification based on SILAC-complemented chemoproteomics.

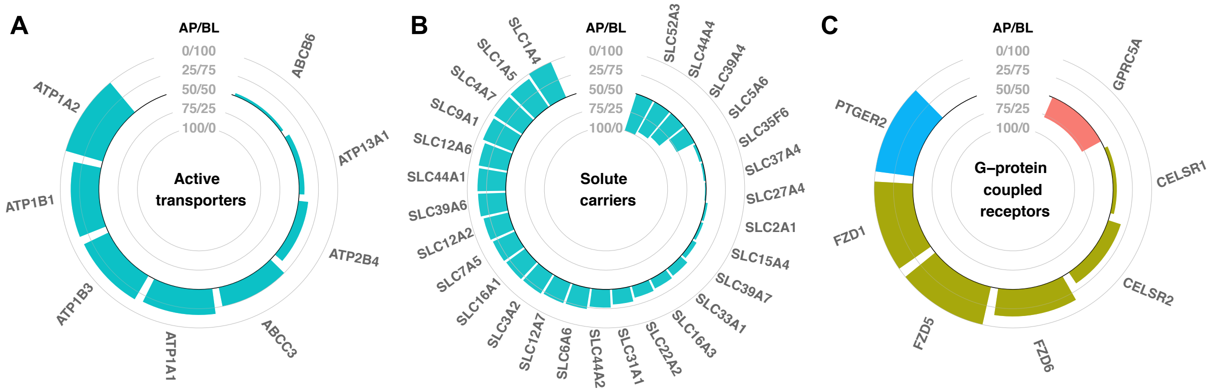


**Figure S2**: Quantitative apicobasal distribution (AP/BL ratio) of selected protein classes on filter-grown MDCK cells based on SILAC-complemented chemoproteomics: **(A)** active transporter proteins, **(B)** solute carrier proteins, **(C)** G-protein coupled receptor proteins. Classification according to the Gene Ontology and KEGG pathway databases (n=4, with 2 SILAC forward and 2 reverse experiments).

**Table S1:** List of all cell surface proteins quantified across the apicobasal surfaceome of filter-grown MDCK cells based on SILAC-complemented chemoproteomics. Apicobasal protein distribution is presented as apical fraction (AP) of the apicobasal protein pool [AP/(AP+BL)] across replicates (fw=SILAC forward; rv=SILAC reverse), and variation is given as median absolute deviation (MAD). Colour-scale indicates descending apical fraction from red to green.

| Gene name | Uniprot ID | AP Median | AP fw1 | AP fw2 | AP rv1 | AP rv2 | MAD |
|---|---|---|---|---|---|---|---|
| ABCB6 | F1PRH1 | 0.465 | 0.377 | 0.462 | 0.469 | 0.495 | 0.031 |
| ABCC3 | J9NY13 | 0.252 | 0.110 | 0.243 | 0.261 | 0.260 | 0.042 |
| ACAT1 | F1PC58 | 0.568 | 0.563 | 0.582 | 0.505 | 0.573 | 0.022 |
| ACE2 | J9P7Y2 | 0.908 | 0.901 | 0.889 | 0.915 | 0.940 | 0.016 |
| ACSF2 | F1PTR3 | 0.470 | 0.521 | 0.477 | 0.436 | 0.463 | 0.025 |
| ADAM10 | F1PGR0 | 0.244 | 0.246 | 0.236 | 0.243 | 0.255 | 0.005 |
| ADAM17 | F1PFZ9 | 0.291 | 0.361 | 0.294 | 0.281 | 0.287 | 0.022 |
| ADAM28 | F1PRX8 | 0.363 | 0.000 | 0.346 | 0.379 | 0.430 | 0.116 |
| ADAM9 | J9P4T2 | 0.389 | 0.409 | 0.417 | 0.352 | 0.370 | 0.026 |
| AGPAT1 | E2RCA0 | 0.497 | 0.519 | 0.495 | 0.467 | 0.498 | 0.014 |
| AGRN | F1Q2Z6 | 0.257 | 0.204 | 0.178 | 0.310 | 0.382 | 0.078 |
| ANO6 | J9NZU1 | 0.193 | 0.112 | 0.174 | 0.236 | 0.212 | 0.040 |
| ANXA2 | Q6TEQ7 | 0.535 | 0.575 | 0.577 | 0.446 | 0.496 | 0.052 |
| APMAP | E2RPE9 | 0.635 | 0.478 | 0.580 | 0.694 | 0.690 | 0.081 |
| ASAH1 | F1PWA9 | 0.557 | 0.416 | 0.557 | NaN | 0.664 | 0.082 |
| ATP13A1 | F1PRS5 | 0.432 | 0.560 | 0.000 | NaN | 0.432 | 0.187 |
| ATP1A1 | J9NZA9 | 0.169 | 0.172 | 0.167 | 0.171 | 0.166 | 0.002 |
| ATP1A2 | F1PL53 | 0.000 | 0.000 | 0.000 | 0.195 | NaN | 0.065 |
| ATP1B1 | P06583 | 0.136 | 0.148 | 0.112 | 0.124 | 0.150 | 0.016 |
| ATP1B3 | F1Q0S2 | 0.146 | 0.136 | 0.156 | 0.110 | 0.216 | 0.032 |
| ATP2B4 | F1PHQ7 | 0.375 | 0.524 | 0.427 | 0.231 | 0.322 | 0.100 |
| B2M | E2RN10 | 0.221 | 0.234 | 0.259 | 0.208 | 0.205 | 0.020 |
| BCAM | J9P3V6 | 0.213 | 0.213 | 0.204 | 0.212 | 0.219 | 0.004 |
| BCHE | F1PEN8 | 0.376 | 0.548 | NaN | NaN | 0.204 | 0.172 |
| BMPR1A | E2QW66 | 0.000 | NaN | NaN | 0.000 | 0.000 | 0.000 |
| BMPR2 | F1PCC1 | 0.000 | NaN | NaN | 0.000 | 0.000 | 0.000 |
| BSG | E2QZT4 | 0.130 | 0.170 | 0.144 | 0.092 | 0.116 | 0.027 |
| BST1 | E2RJT2 | 0.908 | 0.904 | 0.898 | 0.912 | 0.958 | 0.017 |
| BST2 | J9NVI2 | 0.652 | 0.711 | 0.689 | 0.545 | 0.614 | 0.060 |
| CA12 | F1Q440 | 0.243 | 0.220 | 0.141 | 0.266 | 0.338 | 0.061 |
| CA14 | F1P7K9 | 0.369 | 0.383 | 0.369 | 0.369 | 0.256 | 0.032 |
| CADM1 | F1P6Z9 | 0.203 | 0.202 | 0.204 | 0.189 | 0.234 | 0.012 |
| CALU | E2RN38 | 0.894 | 0.474 | 1.000 | 1.000 | 0.788 | 0.184 |
| CANX | P24643 | 0.535 | 0.543 | 0.645 | 0.526 | 0.513 | 0.037 |
| CAV1 | P33724 | 0.538 | 0.600 | 0.564 | 0.490 | 0.512 | 0.041 |
| CAV2 | O46550 | 0.529 | 0.510 | 0.513 | 0.563 | 0.545 | 0.021 |
| CD109 | F1PM26 | 0.586 | 0.451 | 0.393 | 0.757 | 0.721 | 0.158 |
| CD14 (607076) | A0A1S7J0A8 | 1.000 | NaN | NaN | 1.000 | 1.000 | 0.000 |
| CD151 | J9P4E7 | 0.244 | 0.311 | 0.252 | 0.236 | 0.231 | 0.024 |
| CD276 | F1PDP5 | 0.213 | 0.212 | 0.241 | 0.213 | 0.210 | 0.008 |
| CD302 | F1PGD5 | 0.499 | 0.502 | 0.488 | 0.497 | 0.530 | 0.012 |
| CD40 | G1K297 | 0.256 | 0.316 | 0.312 | 0.179 | 0.199 | 0.062 |
| CD44 | J9P423 | 0.217 | 0.213 | 0.214 | 0.220 | 0.265 | 0.014 |
| CD47 | J9P735 | 0.581 | 0.545 | 0.596 | 0.566 | 1.000 | 0.121 |
| CD55 | F1PCI2 | 0.125 | NaN | NaN | 0.249 | 0.000 | 0.125 |
| CD58 | F1PBA0 | 0.097 | 0.047 | 0.074 | 0.121 | 0.135 | 0.034 |
| CD59 | J9P4N4 | 0.732 | 0.738 | 0.773 | 0.697 | 0.726 | 0.022 |
| CD63 | F1PIT4 | 0.507 | 0.699 | 0.718 | 0.314 | 0.250 | 0.213 |
| CD74 | E2RF90 | 0.334 | 0.257 | 0.328 | 0.340 | 0.345 | 0.025 |
| CD81 | F1PB66 | 0.027 | 0.000 | 0.000 | 0.055 | 0.106 | 0.040 |
| CD82 | J9P9V8 | 0.353 | 0.000 | 0.484 | 0.293 | 0.413 | 0.151 |
| CD9 | F1PQM7 | 0.342 | 0.400 | 0.362 | 0.298 | 0.322 | 0.035 |
| CD99 | J9PA14 | 0.258 | 0.253 | 0.263 | 0.267 | 0.250 | 0.007 |
| CDC42 | P60952 | 0.456 | 0.180 | 0.265 | 0.763 | 0.647 | 0.241 |
| CDCP1 | F1PV63 | 0.357 | 0.333 | 0.307 | 0.381 | 0.426 | 0.042 |
| CDH1 | F1PAA4 | 0.196 | 0.195 | 0.171 | 0.219 | 0.197 | 0.013 |
| CDH10 | E2R858 | 0.231 | 0.000 | 0.224 | 0.239 | 0.319 | 0.083 |
| CDH16 | E2REV2 | 0.176 | 0.201 | 0.176 | 0.167 | 0.177 | 0.009 |
| CDH17 | J9NTL4 | 0.521 | 0.469 | 0.457 | 0.573 | 0.576 | 0.056 |
| CDH3 | F1PAA3 | 0.098 | 0.000 | 0.000 | 0.197 | 0.355 | 0.138 |
| CDH6 | E2R0Z0 | 0.418 | 0.413 | 0.402 | 0.428 | 0.424 | 0.009 |
| CEACAM1 | Q004B2 | 0.956 | 0.904 | 0.965 | 0.988 | 0.947 | 0.025 |
| CELSR1 | F1PLY1 | 0.453 | 0.548 | 0.450 | 0.456 | 0.415 | 0.035 |
| CELSR2 | E2R4F0 | 0.329 | 0.350 | 0.331 | 0.286 | 0.327 | 0.017 |
| CENPV | F1PH09 | 0.519 | 0.382 | 0.425 | 0.629 | 0.612 | 0.108 |

| | | | | | | | |
|---|---|---|---|---|---|---|---|
| CEPT1 | E2QV13 | 1.000 | 1.000 | NaN | 0.459 | 1.000 | 0.180 |
| CLDN1 | J9P0K9 | 0.424 | 0.458 | 0.460 | 0.000 | 0.390 | 0.132 |
| CLDN2 | Q95KM6 | 0.400 | 0.483 | 0.446 | 0.301 | 0.353 | 0.069 |
| CLDN3 | Q95KM5 | 0.371 | 0.429 | 0.456 | 0.286 | 0.314 | 0.071 |
| CLDN4 | E2RNC2 | 0.217 | 0.184 | 0.294 | 0.250 | 0.041 | 0.080 |
| CLDN7 | E2R5S3 | 0.101 | 0.202 | 0.000 | NaN | NaN | 0.101 |
| CLDND1 | F1PYH4 | 0.063 | 0.125 | 0.000 | NaN | NaN | 0.063 |
| CLEC7A | E2RFU8 | 0.385 | 0.379 | 0.390 | 0.340 | 0.399 | 0.018 |
| CLU | P25473 | 0.639 | 0.640 | 0.600 | 0.654 | 0.639 | 0.014 |
| CNNM3 | E2RGW9 | 0.000 | 0.000 | 0.000 | NaN | 0.000 | 0.000 |
| CNNM4 | F1PJ05 | 0.143 | 0.189 | 0.140 | 0.118 | 0.145 | 0.019 |
| COL18A1 | F6Y3U7 | 0.176 | 0.105 | 0.084 | 0.344 | 0.247 | 0.101 |
| COL4A1 | Q28271 | 0.424 | 0.266 | 0.237 | 0.618 | 0.581 | 0.174 |
| COL4A2 | F1Q129 | 0.366 | 0.172 | 0.077 | 0.718 | 0.560 | 0.257 |
| CP | F1PAX2 | 0.839 | 0.824 | 0.842 | 0.837 | 0.851 | 0.008 |
| CPM | F1PGL5 | 0.634 | 0.661 | 0.669 | 0.566 | 0.608 | 0.039 |
| CRELD1 | F1PD72 | 0.613 | 0.630 | 0.596 | 0.732 | 0.535 | 0.058 |
| CRIM1 | E2R003 | 0.000 | 0.000 | 0.000 | 0.000 | 0.285 | 0.071 |
| CS | F1PV92 | 0.487 | 0.551 | 0.539 | 0.402 | 0.436 | 0.063 |
| CTNNB1 | B6V8E6 | 0.453 | 0.515 | 0.317 | 0.490 | 0.416 | 0.068 |
| CTSA | F1P9I9 | 0.504 | 0.362 | 0.428 | 0.580 | 0.582 | 0.093 |
| CTSB | E2R6Q7 | 0.464 | 0.506 | 0.357 | 0.426 | 0.502 | 0.056 |
| CTSD | Q4LAL9 | 0.458 | 0.479 | 0.499 | 0.378 | 0.437 | 0.041 |
| CTSH | F6X9C1 | 0.187 | NaN | 0.000 | NaN | 0.374 | 0.187 |
| CTSS | Q8HY81 | 0.489 | NaN | 0.000 | 0.489 | 0.556 | 0.185 |
| CTSZ | F1PIF2 | 0.433 | 0.000 | 0.530 | 0.407 | 0.459 | 0.145 |
| CXADR | F1P992 | 0.296 | 0.325 | 0.289 | 0.266 | 0.302 | 0.018 |
| CYB5R1 | E2RF18 | 0.530 | 0.663 | 0.565 | 0.416 | 0.496 | 0.079 |
| CYP20A1 | F1PLQ3 | 0.495 | 0.493 | 0.490 | 0.498 | 0.545 | 0.015 |
| DAG1 | Q9TSZ6 | 0.375 | 0.340 | 0.325 | 0.503 | 0.411 | 0.062 |
| DAO | E2QV61 | 0.580 | 0.539 | 0.696 | 0.618 | 0.542 | 0.058 |
| DCBLD2 | F1PVX2 | 0.451 | 0.443 | 0.436 | 0.472 | 0.459 | 0.013 |
| DDO | F1PAL6 | 0.512 | 0.531 | 0.599 | 0.426 | 0.493 | 0.053 |
| DDR1 | F1PGJ7 | 0.289 | 0.278 | 0.296 | 0.282 | 0.318 | 0.013 |
| DDX17 | F1PID8 | 0.517 | 0.445 | 0.504 | 0.543 | 0.530 | 0.031 |
| DHCR7 | E2RNQ5 | 0.497 | 0.564 | 0.568 | 0.351 | 0.431 | 0.088 |
| dhrs4 | Q0E9S7 | 0.556 | 0.385 | 0.567 | 0.546 | 0.566 | 0.050 |
| DNAJC10 | E2RCY4 | 0.484 | 0.000 | 0.546 | 0.496 | 0.471 | 0.143 |
| DNASE1L1 | F1PWT3 | 0.670 | 0.684 | 0.694 | 0.592 | 0.655 | 0.033 |
| DPEP1 | E2R4I6 | 1.000 | NaN | 1.000 | 1.000 | 1.000 | 0.000 |
| DSC2 | F1PEF1 | 0.251 | 0.251 | 0.246 | 0.252 | 0.256 | 0.003 |
| DSC3 | F1PEF0 | 0.402 | 0.391 | 0.425 | 0.413 | 0.376 | 0.018 |
| DSG2 | F1PGX2 | 0.476 | 0.518 | 0.502 | 0.450 | 0.438 | 0.033 |
| ECE1 | F1PK66 | 0.000 | 0.000 | 0.000 | 0.203 | 0.000 | 0.051 |
| EFNA1 | E2QUE4 | 0.177 | 0.191 | 0.000 | 0.296 | 0.163 | 0.081 |
| EFNB1 | E2QSB5 | 0.434 | 0.382 | 0.397 | 0.486 | 0.472 | 0.045 |
| EGF | J9NZ75 | 0.533 | 0.553 | 0.792 | 0.506 | 0.512 | 0.082 |
| EGFR | F1PF03 | 0.401 | 0.417 | 0.386 | 0.399 | 0.403 | 0.009 |
| ELOVL1 | E2RGY9 | 0.533 | 0.553 | 0.514 | 0.563 | 0.464 | 0.035 |
| EMC1 | F1PN78 | 0.545 | NaN | 0.618 | 0.545 | 0.219 | 0.133 |
| ENPEP | F6XRM5 | 1.000 | 1.000 | 1.000 | 1.000 | 1.000 | 0.000 |
| ENPP6 | F1PZF9 | 1.000 | 1.000 | 1.000 | 1.000 | 0.835 | 0.041 |
| ENTPD3 | E2RE57 | 0.826 | 0.819 | 0.815 | 0.834 | 0.864 | 0.016 |
| EPCAM | F1PM51 | 0.177 | 0.164 | 0.165 | 0.229 | 0.189 | 0.022 |
| EPHA1 | E2R6A6 | 0.147 | 0.085 | 0.112 | 0.183 | 0.193 | 0.045 |
| EPHA2 | F1Q4C5 | 0.345 | 0.338 | 0.403 | 0.348 | 0.342 | 0.018 |
| EPHB2 | F1PAF1 | 0.357 | 0.380 | 0.329 | 0.406 | 0.333 | 0.031 |
| EPHB4 | F1PRQ8 | 0.358 | 0.383 | 0.357 | 0.358 | 0.308 | 0.019 |
| ERBB2 | F1PIQ9 | 0.255 | 0.256 | 0.236 | 0.253 | 0.267 | 0.009 |
| ERBB3 | F1PVC5 | 0.112 | 0.000 | 0.064 | 0.179 | 0.159 | 0.068 |
| ERBB4 | F1PQ05 | 0.289 | 0.091 | 0.313 | 0.581 | 0.265 | 0.134 |
| ERGIC3 | E2QW07 | 0.644 | NaN | 0.644 | 1.000 | 0.000 | 0.333 |
| ERMP1 | F1PSG1 | 0.505 | 0.521 | 0.489 | 0.545 | 0.433 | 0.036 |
| ERP29 | J9P4L2 | 0.582 | 0.621 | 0.722 | 0.542 | 0.544 | 0.064 |
| ESRP1 | F1PSN8 | 0.383 | 0.383 | 0.511 | 0.303 | NaN | 0.069 |
| F11R | F1PP71 | 0.221 | 0.209 | 0.206 | 0.244 | 0.234 | 0.016 |

| | | | | | | | |
|---|---|---|---|---|---|---|---|
| FAM171A2 | J9P3H5 | 0.182 | 0.233 | 0.073 | 0.131 | 0.272 | 0.075 |
| FAS | F1P7Q7 | 0.209 | 0.285 | 0.133 | 0.063 | 0.328 | 0.104 |
| FCGRT | E2R0L6 | 0.000 | 0.000 | 0.000 | 0.000 | 0.000 | 0.000 |
| FGFR2 | F1PPD8 | 0.270 | 0.293 | 0.127 | 0.264 | 0.276 | 0.045 |
| FGFR3 | F1PK02 | 0.429 | 0.520 | 0.569 | 0.338 | 0.291 | 0.115 |
| FH | E2RGR9 | 0.499 | 0.563 | 0.531 | 0.402 | 0.468 | 0.056 |
| FKBP10 | E2QZA8 | 0.746 | NaN | NaN | 1.000 | 0.492 | 0.254 |
| FLNA | F1PWW0 | 0.427 | 0.283 | 0.376 | 0.537 | 0.478 | 0.089 |
| FLRT2 | E2RB35 | 0.346 | 0.318 | 0.210 | 0.374 | 0.415 | 0.065 |
| FLRT3 | E2RKY7 | 0.451 | 0.445 | 0.360 | 0.459 | 0.456 | 0.028 |
| FLVCR1 | E2RSI2 | 0.305 | 0.352 | 0.368 | 0.258 | 0.175 | 0.072 |
| FN1 | F1P6H7 | 0.458 | 0.316 | 0.386 | 0.531 | 0.791 | 0.155 |
| folh1 | I4IY07 | 0.728 | 0.760 | 0.590 | 0.697 | 0.827 | 0.075 |
| FOLR2 | E2QXF0 | 0.683 | 0.701 | 0.687 | 0.646 | 0.679 | 0.016 |
| FRRS1 | E2RSW3 | 0.267 | NaN | NaN | 0.174 | 0.359 | 0.092 |
| FUCA1 | P48300 | 0.473 | 0.239 | 0.518 | 0.462 | 0.484 | 0.075 |
| FZD1 | F1PCN5 | 0.000 | 0.000 | NaN | 0.000 | NaN | 0.000 |
| FZD5 | E2QVP7 | 0.000 | NaN | 0.000 | 0.000 | 0.000 | 0.000 |
| FZD6 | F1P9V1 | 0.150 | 0.130 | 0.169 | 0.037 | 0.229 | 0.057 |
| GARS | F1Q332 | 0.615 | 0.729 | 0.616 | 0.550 | 0.613 | 0.045 |
| GLG1 | F6V9R9 | 0.537 | 0.522 | 0.474 | 0.555 | 0.552 | 0.028 |
| GLIPR1 | F1PE15 | 0.112 | 0.122 | 0.101 | 0.207 | 0.103 | 0.031 |
| GLS | E2RJ93 | 0.589 | 0.538 | 0.641 | 0.436 | 0.647 | 0.078 |
| GM2A | F1P8U3 | 0.587 | 0.332 | 0.389 | 1.000 | 0.786 | 0.266 |
| GNA13 | F1PZW8 | 0.365 | NaN | NaN | 0.731 | 0.000 | 0.365 |
| GNAI2 | P38400 | 0.461 | 0.461 | 0.515 | 0.418 | 0.460 | 0.025 |
| GNG12 | J9P702 | 0.460 | 0.459 | 0.577 | 0.440 | 0.461 | 0.035 |
| gns | Q32KH4 | 0.549 | NaN | 1.000 | 0.431 | 0.549 | 0.190 |
| GPC1 | F1PPA1 | 0.500 | NaN | 0.753 | 0.415 | 0.500 | 0.113 |
| GPC4 | F1PM35 | 0.083 | 0.000 | 0.000 | 0.257 | 0.165 | 0.106 |
| GPNMB | E2QUR7 | 0.496 | 0.489 | 0.525 | 0.504 | 0.472 | 0.017 |
| GPR107 | F6USA6 | 0.223 | NaN | NaN | 0.000 | 0.445 | 0.223 |
| GPR89A | E2R506 | 0.476 | 0.497 | 0.625 | 0.376 | 0.455 | 0.073 |
| GPRC5A | F1PHN5 | 0.780 | NaN | NaN | 1.000 | 0.561 | 0.220 |
| GRN | E2RHN1 | 0.561 | 0.465 | 0.624 | 0.609 | 0.513 | 0.064 |
| HAVCR1 | F1CME9 | 1.000 | 1.000 | 1.000 | 1.000 | 1.000 | 0.000 |
| HEXB | K0J6C5 | 0.232 | NaN | NaN | 0.000 | 0.463 | 0.232 |
| HFE | F1PX48 | 0.040 | 0.037 | 0.043 | 0.023 | 0.487 | 0.118 |
| HM13 | E2RB97 | 0.554 | 0.554 | 0.555 | NaN | 0.337 | 0.073 |
| HMGCL | E2RFD0 | 0.500 | 0.503 | 0.503 | 0.477 | 0.497 | 0.008 |
| HPN | E2RKP5 | 0.369 | 0.379 | 0.268 | 0.359 | 0.637 | 0.098 |
| HSD17B4 | E2R4T7 | 0.493 | 0.528 | 0.604 | 0.446 | 0.458 | 0.057 |
| HSP90B1 | F1P8N6 | 0.550 | 0.466 | 0.563 | 0.569 | 0.538 | 0.032 |
| HSPG2 | J9NRJ0 | 0.309 | 0.120 | 0.147 | 0.503 | 0.471 | 0.177 |
| HTRA1 | F1PU95 | 0.193 | 0.402 | 0.160 | 0.000 | 0.227 | 0.117 |
| HYAL2 | E2R560 | 0.501 | 0.499 | 0.567 | 0.454 | 0.503 | 0.029 |
| HYOU1 | E2RB31 | 0.498 | 0.528 | 0.638 | 0.468 | 0.394 | 0.076 |
| ICAM1 | P33729 | 0.862 | 0.853 | 0.851 | 0.870 | 0.941 | 0.027 |
| IFI30 | F1PK63 | 0.539 | 0.768 | 0.608 | 0.469 | 0.390 | 0.129 |
| IFNGR1 | F6XBS0 | 0.000 | 0.000 | 0.000 | NaN | 0.000 | 0.000 |
| IGF1R | F1PXU6 | 0.315 | 0.334 | 0.307 | 0.323 | 0.303 | 0.012 |
| IGFBP7 | E2RNL2 | 0.000 | 0.000 | 0.000 | 0.000 | NaN | 0.000 |
| IGSF11 | F1PJL3 | 0.561 | NaN | 0.420 | NaN | 0.702 | 0.141 |
| IGSF3 | F1P9K1 | 0.458 | 0.465 | 0.459 | 0.458 | 0.410 | 0.014 |
| IGSF5 | E2RBD3 | 1.000 | 1.000 | 1.000 | 1.000 | 1.000 | 0.000 |
| IGSF8 | J9NWL5 | 0.409 | NaN | 0.000 | 0.468 | 0.409 | 0.156 |
| IGSF9 | F1PHY9 | 0.325 | 0.347 | 0.333 | 0.316 | 0.293 | 0.018 |
| IL1R1 | F1PSF0 | 0.815 | NaN | 1.000 | 0.815 | 0.779 | 0.074 |
| IL1RAP | J9NU59 | 0.560 | 0.507 | 0.590 | 0.529 | 0.603 | 0.039 |
| IL6ST | F1PTB2 | 0.702 | 0.489 | NaN | 0.702 | 0.950 | 0.154 |
| ITGA1 | F1PP33 | 0.330 | 0.365 | 0.325 | 0.307 | 0.336 | 0.017 |
| ITGA2 | E2REA9 | 0.532 | 0.605 | 0.570 | 0.485 | 0.495 | 0.049 |
| ITGA3 | F1Q439 | 0.250 | 0.283 | 0.240 | 0.230 | 0.259 | 0.018 |
| ITGA6 | E2RL88 | 0.287 | 0.311 | 0.287 | 0.287 | 0.272 | 0.010 |
| ITGAV | F1P8Q0 | 0.429 | 0.460 | 0.446 | 0.412 | 0.401 | 0.023 |
| ITGB1 | E2RT60 | 0.346 | 0.382 | 0.352 | 0.319 | 0.341 | 0.018 |

| | | | | | | | |
|---|---|---|---|---|---|---|---|
| ITGB5 | J9P3C1 | 0.545 | NaN | 0.623 | 0.486 | 0.545 | 0.045 |
| ITGB6 | E2RBL9 | 0.381 | 0.390 | 0.387 | 0.362 | 0.375 | 0.010 |
| ITGB8 | F6X717 | 0.447 | 0.000 | 0.278 | 0.615 | 0.653 | 0.248 |
| ITPR3 | E2RMC0 | 0.473 | 0.372 | 0.445 | 0.525 | 0.502 | 0.053 |
| KIAA1161 | J9PAS5 | 0.076 | 0.087 | 0.000 | 0.076 | NaN | 0.029 |
| KIRREL | F1Q051 | 0.026 | 0.000 | 0.000 | 0.052 | 0.128 | 0.045 |
| KTN1 | E2RG27 | 0.464 | 0.437 | 0.498 | 0.447 | 0.481 | 0.024 |
| L1CAM | E2R2V6 | 0.578 | 0.635 | 0.622 | 0.522 | 0.533 | 0.050 |
| LAMA3 | E2RPP1 | 0.396 | 0.705 | 0.395 | 0.388 | 0.398 | 0.080 |
| LAMB1 | F1PCD8 | 0.422 | 0.456 | 0.238 | 0.447 | 0.397 | 0.067 |
| LAMB3 | F1PFM5 | 0.406 | 0.387 | 0.339 | 0.425 | 0.453 | 0.038 |
| LAMC1 | F1PHK9 | 0.296 | 0.071 | 0.085 | 0.506 | 0.549 | 0.225 |
| LAMP1 | F1Q260 | 0.474 | 0.513 | 0.597 | 0.409 | 0.435 | 0.067 |
| LAMP2 | E2RNJ1 | 0.090 | 1.000 | 0.000 | 0.000 | 0.181 | 0.295 |
| LDLR | F1PML2 | 0.361 | NaN | 0.361 | 0.475 | 0.204 | 0.091 |
| LGALS3BP | E2RKQ6 | 0.539 | 0.562 | 0.594 | 0.516 | 0.499 | 0.035 |
| LGMN | E2QXF2 | 0.485 | 0.481 | 0.310 | 0.489 | 0.567 | 0.066 |
| LHFPL2 | J9P9A6 | 0.516 | 0.524 | 0.533 | 0.394 | 0.507 | 0.039 |
| LIPH | F1PG57 | 0.231 | 0.168 | 0.253 | 0.210 | 0.324 | 0.050 |
| LMAN2 | P49256 | 0.543 | 0.549 | 0.536 | 0.559 | 0.380 | 0.048 |
| LMF2 | J9NVV3 | 0.518 | 0.533 | 0.549 | 0.475 | 0.503 | 0.026 |
| LRP1 | J9P315 | 0.239 | 0.197 | 0.161 | 0.281 | 0.299 | 0.056 |
| LRPAP1 | F1PAG7 | 0.852 | NaN | NaN | 1.000 | 0.705 | 0.148 |
| LSAMP | F1PJN2 | 0.725 | 0.751 | 0.734 | 0.716 | 0.709 | 0.015 |
| LSR | F1PKT2 | 0.130 | 0.115 | 0.096 | 0.148 | 0.145 | 0.020 |
| LTBR | E2RG08 | 0.010 | 0.000 | 0.007 | 0.013 | 0.098 | 0.026 |
| M6PR | E2R4C1 | 0.201 | 0.209 | 0.154 | 0.193 | 0.228 | 0.022 |
| MAGT1 | E2RRH2 | 0.546 | 0.526 | 0.566 | 0.196 | 0.576 | 0.105 |
| MET | Q75ZY9 | 0.406 | 0.403 | 0.360 | 0.416 | 0.409 | 0.015 |
| METTL7A | J9NTA1 | 0.454 | 0.446 | 0.449 | 0.498 | 0.460 | 0.016 |
| MFGE8 | F1PFZ5 | 0.731 | 0.710 | 0.673 | 0.778 | 0.752 | 0.037 |
| MFSD6 | E2RK97 | 0.044 | 0.000 | 0.000 | 0.307 | 0.087 | 0.099 |
| MLEC | E2RD92 | 0.457 | 0.479 | 0.611 | 0.412 | 0.436 | 0.060 |
| MME | F5C3N2 | 1.000 | NaN | NaN | 1.000 | 1.000 | 0.000 |
| MMP14 | F1PXM2 | 0.264 | 0.324 | 0.096 | 0.305 | 0.224 | 0.077 |
| MMP15 | F1P625 | 0.116 | 0.034 | 0.065 | 0.167 | 0.269 | 0.085 |
| MPZL1 | J9NSZ7 | 0.124 | NaN | 0.000 | 0.124 | 0.160 | 0.053 |
| MRPL21 | J9P2W1 | 0.000 | 0.000 | 0.000 | NaN | 0.437 | 0.146 |
| MRPS35 | E2R2D2 | 0.309 | 0.518 | 0.000 | 0.281 | 0.338 | 0.143 |
| MSLN | J9NX38 | 0.834 | 0.977 | 0.881 | 0.654 | 0.787 | 0.104 |
| MUC1 | J9P6J1 | 0.901 | 0.894 | 0.880 | 0.907 | 0.931 | 0.016 |
| MUC20 | F1PJ84 | 0.929 | 0.840 | 0.858 | 1.000 | 1.000 | 0.076 |
| NCLN | E2QY93 | 0.413 | 0.513 | 0.604 | 0.313 | 0.000 | 0.201 |
| NCSTN | M1VEJ1 | 0.267 | 0.249 | 0.000 | 0.285 | 0.394 | 0.107 |
| NDUFA10 | E2RIG0 | 0.489 | 0.500 | 0.418 | 0.477 | 0.598 | 0.051 |
| NEO1 | J9P2T1 | 0.335 | 0.101 | 0.333 | 0.338 | 0.433 | 0.084 |
| NNT | F1PLG2 | 0.467 | 0.547 | 0.491 | 0.295 | 0.443 | 0.075 |
| NOTCH2 | F1PPP5 | 0.226 | 0.000 | 0.214 | 0.298 | 0.238 | 0.081 |
| NPC1L1 | A0MJA4 | 0.929 | 0.934 | 0.931 | 0.927 | 0.923 | 0.004 |
| NPC2 | Q28895 | 0.375 | 0.375 | 0.310 | NaN | 0.679 | 0.123 |
| NPDC1 | F6XGX1 | 0.000 | NaN | 0.000 | NaN | 0.000 | 0.000 |
| NPTN | F1PDQ0 | 0.040 | 0.037 | 0.022 | 0.069 | 0.044 | 0.014 |
| NTN4 | E2QZY0 | 0.180 | 0.000 | NaN | 0.359 | NaN | 0.180 |
| NUP155 | F6XTD9 | 0.411 | 0.256 | 0.437 | 0.384 | 0.460 | 0.064 |
| NUP210 | F1PSK9 | 0.507 | 0.490 | 0.636 | 0.507 | NaN | 0.049 |
| OCLN | E2R3X4 | 0.156 | 0.000 | 0.152 | 0.160 | 0.169 | 0.044 |
| OSMR | E2QWS7 | 0.631 | 0.593 | 0.628 | 0.634 | 0.908 | 0.080 |
| P4HA1 | E2RLB8 | 0.591 | 0.606 | 0.646 | 0.575 | 0.504 | 0.043 |
| P4HA2 | F1PTP6 | 0.552 | 0.649 | NaN | NaN | 0.455 | 0.097 |
| PCCB | F1PTU4 | 0.482 | 0.533 | 0.508 | 0.406 | 0.457 | 0.045 |
| PCDH1 | E2QX93 | 0.219 | 0.191 | 0.151 | 0.247 | 0.248 | 0.038 |
| PCYOX1 | E2R1N7 | 0.388 | 0.243 | 1.000 | 0.534 | 0.203 | 0.272 |
| PGAP1 | E2R8V9 | 0.493 | 0.502 | NaN | 0.493 | 0.488 | 0.005 |
| PIGN | F1PVD7 | 0.689 | 1.000 | NaN | NaN | 0.379 | 0.311 |
| PIGO | E2RKS8 | 0.494 | 0.557 | 0.516 | 0.473 | 0.436 | 0.041 |
| PIGR | F6Y6T8 | 0.411 | 0.369 | 0.316 | 0.452 | 0.496 | 0.066 |

| | | | | | | | |
|---|---|---|---|---|---|---|---|
| PLAUR | E2RGN0 | 0.804 | 0.795 | 0.666 | 0.814 | 0.831 | 0.046 |
| PLD3 | E2QVP0 | 0.350 | NaN | 0.561 | 0.350 | 0.339 | 0.074 |
| PLOD2 | F1PZL5 | 0.448 | 0.407 | 0.337 | 0.857 | 0.489 | 0.150 |
| PLOD3 | E2RTL4 | 0.389 | NaN | 0.000 | 0.389 | 1.000 | 0.333 |
| PLXNA1 | F1P6G0 | 0.216 | 0.141 | 0.178 | 0.255 | 0.392 | 0.082 |
| PLXNA2 | F1PHZ1 | 0.421 | 0.441 | 0.447 | 0.400 | 0.327 | 0.040 |
| PLXNB1 | E2RM56 | 0.213 | 0.215 | 0.215 | 0.209 | 0.212 | 0.003 |
| PLXNB2 | F1P9S5 | 0.307 | 0.320 | 0.309 | 0.292 | 0.304 | 0.008 |
| PLXNB3 | F1PXM6 | 0.000 | 0.000 | 0.209 | NaN | 0.000 | 0.070 |
| PMPCA | F1PF09 | 0.488 | 0.513 | 0.463 | 0.407 | 0.632 | 0.069 |
| PNPLA6 | E2RIR2 | 0.497 | 0.379 | 0.504 | 0.497 | NaN | 0.042 |
| PODXL | F1PID4 | 0.928 | 0.917 | 0.918 | 0.937 | 0.950 | 0.013 |
| PON2 | F1PSG5 | 0.483 | 0.403 | 0.513 | 0.454 | 0.512 | 0.042 |
| PPIB | F1PLV2 | 0.703 | 0.555 | 0.754 | 0.741 | 0.666 | 0.069 |
| PRCP | F1PWK3 | 0.264 | 0.000 | NaN | NaN | 0.528 | 0.264 |
| PRKCSH | E2RKK6 | 0.795 | 0.590 | 1.000 | 1.000 | 0.579 | 0.208 |
| PrP | O46593 | 0.697 | 0.698 | 0.707 | 0.656 | 0.697 | 0.013 |
| PTGER2 | F1P776 | 0.000 | 0.000 | 0.000 | NaN | NaN | 0.000 |
| PTGFRN | F1PR26 | 0.346 | 0.379 | 0.370 | 0.322 | 0.322 | 0.026 |
| PTK7 | E2RMT2 | 0.433 | 0.462 | 0.423 | 0.395 | 0.443 | 0.022 |
| PTPRA | E2R5J7 | 0.000 | NaN | 0.000 | NaN | 0.000 | 0.000 |
| PTPRD | F1PH17 | 0.492 | 0.441 | 0.413 | 0.564 | 0.542 | 0.063 |
| PTPRF | E2RE80 | 0.364 | 0.337 | 0.334 | 0.402 | 0.391 | 0.030 |
| PTPRG | F1PWC1 | 0.272 | 0.184 | 0.259 | 0.288 | 0.285 | 0.033 |
| PTPRJ | F1P983 | 0.941 | 0.971 | 0.867 | 0.931 | 0.951 | 0.031 |
| PTPRM | F1PUP2 | 0.422 | 0.421 | 0.431 | 0.403 | 0.423 | 0.008 |
| PTPRO | F1PWH3 | 1.000 | 0.744 | 1.000 | 1.000 | 1.000 | 0.064 |
| PTPRS | E2R6Z2 | 0.154 | 0.078 | 0.065 | 0.230 | 0.310 | 0.099 |
| PVR | F1PVS7 | 0.362 | 0.390 | 0.384 | 0.340 | 0.309 | 0.031 |
| RAB10 | F2Z4P9 | 0.464 | 0.553 | 0.489 | 0.410 | 0.440 | 0.048 |
| RAB13 | F1PTE3 | 0.459 | 0.476 | 0.000 | 0.493 | 0.443 | 0.131 |
| RAB25 | E2RQ15 | 0.500 | 0.533 | 0.581 | 0.466 | 0.437 | 0.053 |
| RAB5A | P18066 | 0.456 | 0.404 | 0.499 | 0.443 | 0.468 | 0.030 |
| RAB5C | P51147 | 0.470 | 0.375 | 0.467 | 0.541 | 0.473 | 0.043 |
| RAB8A | P61007 | 0.460 | 0.502 | 0.512 | 0.407 | 0.418 | 0.047 |
| RAB9A | P24408 | 0.525 | NaN | 0.381 | 0.598 | 0.525 | 0.072 |
| RAC1 | P62999 | 0.482 | 0.474 | 0.490 | 0.389 | 0.536 | 0.041 |
| RDH14 | J9NWS8 | 0.485 | 0.381 | 0.511 | 0.459 | 0.594 | 0.066 |
| RELL1 | E2QUZ0 | 0.129 | 0.149 | 0.113 | 0.027 | 0.146 | 0.039 |
| RPN1 | E2RQ08 | 0.515 | 0.538 | 0.563 | 0.478 | 0.493 | 0.033 |
| RPSA | J9JHJ2 | 0.421 | 0.422 | 0.420 | 0.412 | 0.520 | 0.027 |
| RTN4 | E2R925 | 0.582 | 0.565 | 0.599 | 0.624 | 0.553 | 0.026 |
| SCARB1 | F1PQT1 | 1.000 | 1.000 | 1.000 | NaN | NaN | 0.000 |
| SCPEP1 | J9NRZ7 | 0.536 | 0.618 | 0.453 | 1.000 | 0.096 | 0.267 |
| SDC4 | J9NZ79 | 0.082 | 0.044 | 0.041 | 0.126 | 0.121 | 0.040 |
| SDHB | Q0QEY4 | 0.492 | 0.543 | 0.682 | 0.362 | 0.441 | 0.106 |
| SEC61A1 | J9NXA9 | 0.487 | 0.581 | 0.492 | 0.479 | 0.482 | 0.028 |
| SEMA4B | F1P9X6 | 0.311 | 0.203 | 0.230 | 0.411 | 0.392 | 0.092 |
| SEMA4C | E2RHV1 | 0.356 | 0.271 | 0.195 | 0.440 | 0.453 | 0.107 |
| SEMA4D | F1PSB1 | 0.381 | 0.315 | 0.299 | 0.447 | 0.485 | 0.079 |
| SEMA4G | E2RP18 | 0.104 | 0.000 | 0.038 | 0.199 | 0.170 | 0.083 |
| SERPINH1 | E2RHY7 | 0.497 | 0.481 | 0.544 | 0.485 | 0.508 | 0.022 |
| SFTPD | E2QWR0 | 0.683 | NaN | 0.000 | 0.683 | 1.000 | 0.333 |
| SIAE | F1PXN3 | 0.531 | 0.454 | 0.609 | NaN | NaN | 0.077 |
| SIGIRR | F1PY07 | 0.083 | 0.000 | 0.000 | 0.166 | 0.172 | 0.085 |
| SLC12A2 | F1P9W5 | 0.167 | 0.107 | 0.130 | 0.203 | 0.281 | 0.062 |
| SLC12A6 | F1Q2J3 | 0.120 | 0.127 | 0.416 | 0.000 | 0.113 | 0.107 |
| SLC12A7 | F1PKH1 | 0.220 | 0.000 | 0.354 | 0.220 | NaN | 0.118 |
| SLC15A4 | F1PWH8 | 0.463 | 0.619 | NaN | 0.463 | 0.444 | 0.058 |
| SLC16A1 | F1PCQ3 | 0.195 | 0.218 | 0.173 | 0.180 | 0.209 | 0.018 |
| SLC16A3 | J9NVM8 | 0.363 | 0.405 | 0.456 | 0.249 | 0.322 | 0.072 |
| SLC1A4 | F1Q0H9 | 0.000 | 0.000 | 0.000 | 0.000 | 0.000 | 0.000 |
| SLC1A5 | K7ZSN9 | 0.053 | 0.000 | 0.059 | 0.047 | 0.307 | 0.080 |
| SLC22A2 | F1Q431 | 0.344 | 0.426 | NaN | 0.261 | NaN | 0.082 |
| SLC27A4 | F1PAM6 | 0.515 | 0.514 | 0.516 | 0.471 | 1.000 | 0.132 |
| SLC2A1 | F1PWN2 | 0.468 | 0.506 | 0.553 | 0.346 | 0.430 | 0.071 |

| | | | | | | | |
|---|---|---|---|---|---|---|---|
| SLC31A1 | J9P9D2 | 0.302 | 0.000 | 0.405 | 0.308 | 0.295 | 0.104 |
| SLC33A1 | E2RTD2 | 0.380 | 0.356 | 0.583 | 0.404 | 0.335 | 0.074 |
| SLC35F6 | J9P2U5 | 0.537 | 0.536 | 0.538 | 0.639 | 0.444 | 0.049 |
| SLC37A4 | F1PPG9 | 0.527 | 0.588 | 0.427 | 0.544 | 0.509 | 0.049 |
| SLC39A4 | F1PC23 | 0.908 | 0.854 | 0.901 | 0.929 | 0.914 | 0.022 |
| SLC39A6 | E2RMA6 | 0.139 | 0.177 | 0.015 | 0.285 | 0.101 | 0.086 |
| SLC39A7 | Q5TJF6 | 0.435 | 0.501 | 0.000 | 0.435 | NaN | 0.167 |
| SLC3A2 | F1PRC5 | 0.202 | 0.233 | 0.219 | 0.185 | 0.171 | 0.024 |
| SLC44A1 | E2R949 | 0.133 | 0.134 | 0.099 | 0.131 | 0.139 | 0.011 |
| SLC44A2 | J9P0D9 | 0.264 | 0.281 | 0.219 | 0.246 | 0.352 | 0.042 |
| SLC44A4 | F1P9U5 | 0.947 | 0.880 | 0.954 | 0.950 | 0.944 | 0.020 |
| SLC4A7 | J9PAF7 | 0.063 | 0.060 | 0.082 | 0.042 | 0.066 | 0.012 |
| SLC52A3 | E2RM06 | 1.000 | NaN | 1.000 | 1.000 | 1.000 | 0.000 |
| SLC5A6 | E2R9T1 | 0.763 | 0.911 | 0.726 | 0.801 | 0.689 | 0.074 |
| SLC6A6 | Q00589 | 0.226 | 0.296 | 0.000 | 0.175 | 0.276 | 0.099 |
| SLC7A5 | F1PH98 | 0.183 | 0.264 | 0.172 | 0.186 | 0.180 | 0.025 |
| SLC9A1 | F1PID1 | 0.079 | 0.121 | 0.066 | 0.054 | 0.091 | 0.023 |
| SLIT3 | F1Q3G2 | 0.381 | 0.197 | 0.128 | 0.564 | 0.567 | 0.202 |
| SLITRK4 | F1Q4K7 | 0.417 | 0.390 | 0.439 | 0.402 | 0.431 | 0.020 |
| SPINT1 | F1PQ47 | 0.354 | 0.334 | 0.328 | 0.384 | 0.374 | 0.024 |
| SPINT2 | F1P984 | 0.225 | 0.173 | 0.201 | 0.249 | 0.251 | 0.032 |
| SSR1 | P16967 | 0.512 | 0.478 | 0.546 | 0.139 | 0.713 | 0.160 |
| SSR4 | E2QY54 | 0.544 | 0.589 | 0.635 | 0.452 | 0.500 | 0.068 |
| ST14 | F1PF63 | 0.206 | 0.187 | 0.173 | 0.225 | 0.311 | 0.044 |
| STEAP3 | F1P8R0 | 0.288 | 0.240 | 0.277 | 0.298 | 0.416 | 0.049 |
| STT3A | F1PJP5 | 0.508 | 0.510 | 0.548 | 0.480 | 0.507 | 0.017 |
| STT3B | E2RG47 | 0.476 | 0.548 | 0.541 | 0.299 | 0.412 | 0.094 |
| SUCLG1 | E2R0Y5 | 0.444 | 0.418 | 0.544 | 0.313 | 0.469 | 0.071 |
| SUOX | F1PVD4 | 0.500 | 0.275 | 0.441 | 1.000 | 0.560 | 0.211 |
| SYNGR2 | F1PZ37 | 1.000 | 1.000 | 1.000 | NaN | 0.000 | 0.333 |
| SYPL1 | E2RC10 | 0.324 | 0.327 | 0.337 | 0.242 | 0.320 | 0.025 |
| TAPBP | Q5TJE4 | 0.535 | 0.598 | 0.594 | 0.477 | 0.472 | 0.061 |
| TBL2 | E2RP67 | 0.478 | 0.532 | 0.566 | 0.346 | 0.424 | 0.082 |
| TENM4 | F1PFD6 | 0.502 | 0.655 | 0.543 | 0.461 | 0.439 | 0.074 |
| TEX264 | E2RAC8 | 0.464 | NaN | 0.552 | NaN | 0.376 | 0.088 |
| TF | F6V1W9 | 0.438 | 0.437 | 0.506 | 0.438 | 0.437 | 0.018 |
| TGFBR3 | F1PIG0 | 0.310 | 0.310 | 0.311 | 0.299 | 0.352 | 0.014 |
| TIMM21 | E2R117 | 0.496 | NaN | 0.627 | 0.000 | 0.496 | 0.209 |
| TINAGL1 | E2QXH3 | 0.249 | 0.000 | 0.187 | 0.330 | 0.310 | 0.113 |
| TM2D3 | F6UUM9 | 0.234 | 0.606 | NaN | 0.234 | 0.177 | 0.143 |
| TM7SF2 | E2RS95 | 0.512 | 0.565 | 0.575 | 0.444 | 0.460 | 0.059 |
| TM9SF2 | E2R5Z5 | 0.416 | 0.513 | 0.427 | 0.232 | 0.404 | 0.076 |
| TM9SF3 | E2RSD0 | 0.441 | 0.571 | 0.474 | 0.290 | 0.407 | 0.087 |
| TMED10 | E2QYL0 | 0.537 | 0.581 | 0.698 | 0.493 | 0.464 | 0.080 |
| TMED9 | F6XD36 | 0.495 | 0.609 | 0.529 | 0.458 | 0.461 | 0.055 |
| TMEM109 | J9P616 | 0.605 | 0.547 | 0.662 | 0.793 | 0.471 | 0.109 |
| TMEM2 | F1PH14 | 0.353 | 0.323 | 0.329 | 0.399 | 0.378 | 0.031 |
| TMEM30A | E2R2Z8 | 0.349 | 0.437 | 0.418 | 0.067 | 0.280 | 0.127 |
| TMPRSS11E | F1Q3A2 | 0.297 | 0.319 | 0.303 | 0.270 | 0.290 | 0.015 |
| TMX1 | E2RK67 | 0.533 | 0.590 | 0.560 | 0.434 | 0.506 | 0.052 |
| TMX3 | F1PIX5 | 0.516 | 0.432 | NaN | 0.516 | 0.681 | 0.083 |
| TPBG | E2QXI6 | 0.293 | 0.230 | 0.287 | 0.336 | 0.298 | 0.029 |
| TRPA1 | F1Q2M0 | 0.000 | 0.000 | 0.414 | 0.000 | 0.000 | 0.104 |
| TSPAN1 | J9NS67 | 0.645 | 0.537 | NaN | 0.645 | 1.000 | 0.154 |
| TSPAN14 | E2QZH1 | 0.228 | 0.113 | 0.214 | 0.251 | 0.241 | 0.042 |
| TSPAN15 | E2RS68 | 0.128 | 0.054 | 0.127 | 0.129 | 0.167 | 0.029 |
| TSPAN33 | F1PEJ5 | 0.034 | 0.000 | 0.000 | 0.067 | 0.149 | 0.054 |
| TSPAN6 | F1PMP0 | 0.000 | 0.000 | 0.000 | 0.000 | NaN | 0.000 |
| TSPAN8 | E2QSC8 | 0.220 | 0.236 | 0.240 | 0.190 | 0.204 | 0.020 |
| TTYH3 | F1P9A7 | 0.317 | 0.335 | 0.134 | 0.302 | 0.332 | 0.058 |
| UNC93B1 | F1PXK2 | 0.414 | 0.549 | 0.000 | 0.348 | 0.480 | 0.170 |
| UPK3B | E2R4D0 | 0.838 | 0.830 | 0.861 | 0.846 | 0.829 | 0.012 |
| VASN | E2RRN8 | 0.121 | 0.051 | 0.077 | 0.165 | 0.165 | 0.051 |
| VASP | P50551 | 0.524 | 0.452 | 0.524 | 0.524 | 0.548 | 0.024 |
| VCAN | J9P884 | 0.197 | 0.053 | 0.138 | 0.255 | 0.279 | 0.086 |
| VDAC2 | E2R948 | 0.476 | 0.576 | 0.541 | 0.344 | 0.412 | 0.091 |

| | | | | | | | |
|---|---|---|---|---|---|---|---|
| VLDLR | F1P787 | 0.100 | 0.119 | 0.082 | 0.039 | 0.300 | 0.074 |
| VNN1 | Q9TSX8 | 0.750 | 0.736 | 0.780 | 0.738 | 0.762 | 0.017 |
| VSIG10 | J9P1R7 | 0.176 | 0.241 | 0.000 | 1.000 | 0.111 | 0.282 |
| WFDC2 | F1PDT8 | 0.486 | 0.566 | 0.486 | 0.358 | NaN | 0.069 |
